# AscA (YecA) is a molecular chaperone involved in Sec-dependent protein translocation in *Escherichia coli*

**DOI:** 10.1101/2020.07.21.215244

**Authors:** Tamar Cranford Smith, Max Wynne, Cailean Carter, Chen Jiang, Mohammed Jamshad, Mathew T. Milner, Yousra Djouider, Emily Hutchinson, Peter A. Lund, Ian Henderson, Damon Huber

## Abstract

Proteins that are translocated across the cytoplasmic membrane by Sec machinery must be in an unfolded conformation in order to pass through the protein-conducting channel during translocation. Molecular chaperones assist Sec-dependent protein translocation by holding substrate proteins in an unfolded conformation in the cytoplasm until they can be delivered to the membrane-embedded Sec machinery. For example, in *Escherichia coli*, SecB binds to a subset of unfolded Sec substrates and delivers them to the Sec machinery by interacting with the metal-binding domain (MBD) of SecA, an ATPase required for translocation in bacteria. Here, we describe a novel molecular chaperone involved Sec-dependent protein translocation, which we have named AscA (for <u>a</u>ccessory <u>S</u>ec <u>c</u>omponent). AscA contains a metal-binding domain (MBD) that is nearly identical to the MBD of SecA. *In vitro* binding studies indicated that AscA binds to SecB and ribosomes in an MBD-dependent fashion.

Saturated transposon mutagenesis and genetics studies suggested that AscA is involved in cell-envelope biogenesis and that its function overlaps with that of SecB. In support of this idea, AscA copurified with a range of proteins and prevented the aggregation of citrate synthase *in vitro*. Our results suggest that AscA is molecular chaperone and that it enhances Sec-dependent protein translocation by delivering its substrate proteins to SecB.

**IMPORTANCE:** This research describes the discovery of a novel molecular chaperone, AscA (YecA). The function of AscA was previously unknown. However, it contains a small domain, known as the MBD, suggesting it could interact with the bacterial Sec machinery, which is responsible for transporting proteins across the cytoplasmic membrane. The work described this study indicates that the MBD allows AscA to bind to both the protein synthesis machinery and the Sec machinery. The previously function of the previously uncharacterised N-terminal domain is that of a molecular chaperone, which binds to unfolded substrate proteins. We propose that AscA binds to protein substrates as they are still be synthesised by ribosomes in order to channel them into the Sec pathway.

## INTRODUCTION

In *Escherichia coli*, most newly synthesized periplasmic and outer membrane proteins are transported across the cytoplasmic membrane by the Sec machinery. During translocation, protein substrates of the Sec machinery pass through an evolutionarily conserved channel in the cytoplasmic membrane (composed of the integral membrane proteins SecY, -E and -G) in an unfolded conformation (1, 2). In addition, translocation usually requires the activity of SecA (3), an ATPase that facilitates translocation through SecYEG (4). The translocation of periplasmic and outer membrane proteins typically begins only after the substrate protein is fully (or nearly fully) synthesised (*i.e.* “posttranslationally”) (5, 6).

Because proteins must be unfolded to pass through SecYEG, folding of substrate proteins in the cytoplasm blocks Sec-dependent protein translocation, causing a protein to become irreversibly trapped in the cytoplasm (7). Furthermore, partially folded proteins that engage SecYEG can clog (or “jam”) the Sec machinery, which is toxic (8). As a result, cells have evolved multiple mechanisms to prevent premature folding of substrate proteins. For example, molecular chaperones can bind to unfolded Sec substrate proteins and hold them in an unfolded conformation until they can be delivered to the membrane-embedded Sec machinery. One such chaperone is SecB, which binds to a subset of unfolded Sec substrate proteins and delivers them to SecA for translocation across the membrane (9–13). Recognition of nascent substrates by SecB is dependent on SecA (14), suggesting that SecB requires an intermediary to recognise its substrate proteins.

The interaction of SecA with SecB is mediated by a small (∼20 amino acid) metal-binding domain (MBD) near the extreme C-terminus of SecA (13, 15, 16). Recent work indicates that the MBD also binds to ribosomes and that ribosome binding is involved in coordinating binding of SecA to nascent polypeptides (17). As its name indicates, the MBD binds to a transition metal (zinc and/or iron) (15, 18), and binding to the metal ion is required for stable folding of the MBD (15). The amino acids responsible for metal binding are highly conserved (C*X*C*X*S*X_6_*CH or C*X*C*X*S*X_6_*CC) (15, 17, 19).

We recently described a protein of unknown function in *E. coli* that contains a MBD that is nearly identical to the MBD of SecA (18), YecA, which we have re-named AscA (for <u>a</u>ccessory <u>S</u>ec <u>c</u>omponent). AscA also contains a UPF0149-family domain at its N-terminus, the function of which has not been described. In this work, we investigated the function of AscA. The similarity of the AscA and SecA MBDs led us to investigate the interaction of AscA with SecB and ribosomes and the dependence of these interactions on the MBD. Genetic analysis suggested that AscA is involved in cell-envelope biogenesis and that AscA could be a molecular chaperone. Further studies indicated that AscA binds to cytoplasmic Sec substrate proteins and that it carries out its function in coordination with SecB *in vivo*. Our results suggest a potential model for how AscA could facilitate Sec-dependent protein translocation in *E. coli*.

## RESULTS

### Binding of AscA to SecB

Many of the amino acids that mediate the interaction between the SecA MBD and SecB from *Haemophilus influenzae* are conserved in the MBD of AscA (**supplemental figure S1**) (18, 20). To investigate whether AscA can also bind to SecB, we examined the effect of AscA on the thermophoretic mobility of SecB using microscale thermophoresis. To this end, we fluorescently labelled SecB and incubated it with unlabelled AscA. There was a large change in the thermophoretic properties of fluorescently labelled SecB at saturating concentrations of AscA (**figure 1A**), suggesting that SecB binds to AscA. However, the presence of a truncated variant of AscA, which lacks the MBD (AscAΔMBD), did not affect thermophoresis of SecB (**figure 1A**). Purified AscAΔMBD was fully folded even in the absence of its MBD (18), indicating that the interaction between SecB and AscA is dependent on the MBD. Analysis of the effect of increasing concentrations of AscA on the thermophoresis of suggested an equilibrium dissociation constant (K_D_) of approximately 150 nM (**figure 1B**).

**Figure 1.**
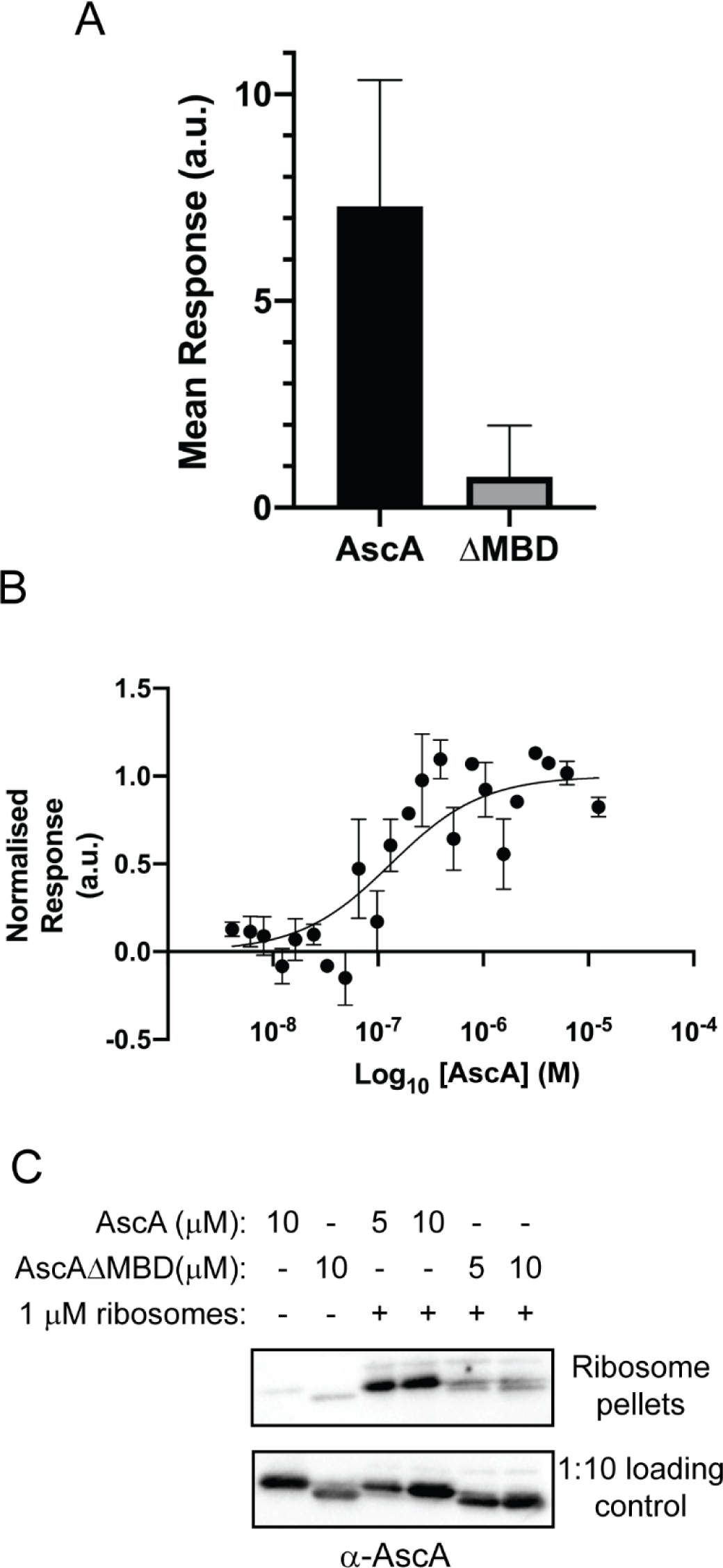
Binding of AscA to SecB and to ribosomes is dependent on its C-terminal MBD. (A & B) Fluorescently labelled SecB was incubated in the absence or presence of AscA or AscAΔMBD, and the effect of AscA on the thermophoretic mobility of SecB was determined by microscale thermophoresis. (A) The magnitude of the effect of 100 μM unlabelled AscA or AscAΔMBD on the thermophoretic mobility of 160 nM SecB was determined by microscale thermophoresis. Confidence intervals represent one SD. (B) To determine the approximate dissociation constant of the AscA-SecB complex, 160 nM fluorescently labelled SecB was incubated in the presence of AscA at concentrations between 6 nM and 200 μM, and the K_D_ was determined by fitting the data. Thermophoresis was determined at least three time for each AscA concentration. Confidence intervals are one standard deviation. (C) The indicated concentrations of AscA or AscAΔMBD were incubated in the absence or presence of 1 μM vacant 70S ribosomes. After equilibration at 30°C, the binding reactions were layered on a 30% sucrose cushion and centrifuged at >200,000 x *g*. The pellet fractions were resolved by SDS-PAGE and subjected to western blotting using anti-AscA antiserum (above). As a loading control, a 1:10 dilution of the binding reaction prior to centrifugation was also analysed by western blotting (below).

### Binding of AscA to ribosomes

We next investigated the interaction of AscA with ribosomes. To this end, we incubated AscA or AscAΔMBD with purified non-translating 70S ribosomes and then separated ribosome-bound AscA from unbound AscA by sedimenting ribosomes through a 30% sucrose cushion by ultracentrifugation. Full-length AscA cosedimented with the 70S ribosomes, indicating that it can bind to ribosomes (**figure 1C**).

Truncation of the MBD in AscAΔMBD greatly reduced its ability to cosediment with ribosomes, indicating that binding is dependent on the MBD. Ribosome cosedimentation experiments in the presence of increasing concentrations of AscA suggested that the K_D_ of the AscA-ribosome complex was in the μM range and that binding saturated at a 1:1 stoichiometry (**supplemental figure S2**).

### Transposon-directed insertion-site sequencing of a Δ<u>ascA</u> mutant

To investigate the function of AscA, we determined which genes are essential for viability in a Δ*ascA* deletion mutant using <u>tra</u>nsposon-<u>d</u>irected insertion-site <u>s</u>equencing (TraDIS) (21, 22). Briefly, we created a library of ∼500,000 independent mini-Tn*5* transposon insertion mutants in a Δ*ascA* deletion mutant (23) (BW25113 Δ*ascA*) and then determined the location of each transposon insertion site using Illumina sequencing. Analysis of the location of the insertion sites revealed a set of ∼143 genes that did not contain any insertions in BW25113 Δ*ascA* but did contain insertions in the isogenic parent (**supplemental table S1**). These mutations affected a range of process but predominantly affected cell envelope biogenesis, cell-envelope stress responses, cell division, protein folding and protein synthesis. These results suggest that the effect of the Δ*ascA* mutation is pleiotropic, consistent with a role in cell envelope biogenesis.

### Expression of AscA

The *ascA* gene is co-transcribed in a polycistronic message, which contains binding sites for the sRNAs RyhB and RybB immediately 5′ to the *ascA* cistron (24). RyhB typically represses genes encoding iron-containing proteins, and RybB is involved in the σ^E^ cell-envelope stress response, suggesting that expression of AscA could be dependent on iron availability or cell envelope stress. To investigate this possibility, we examined the steady-state level of AscA in *E. coli* using by western blotting. Initial experiments indicated that the presence of ampicillin in the growth medium increased the steady state levels of AscA in the cells, suggesting that expression is dependent on cell envelope stress (**supplemental figure S3A**). Subinhibitory concentrations of chloramphenicol did not induce AscA expression (data not shown). In addition, BW25113 expressed AscA at higher levels when grown at lower temperatures (**figure S3B**). Finally, the presence of EDTA (which chelates transition metal ions with high affinity) completely repressed expression of AscA (**supplemental figure S3C**), consistent with the regulation of AscA by RyhB. These results suggest that expression of AscA is regulated by cell envelope stress and iron availability.

### Construction of a Δ<u>ascA</u> Δ<u>secB</u> double mutant

Because AscA and SecB physically interact with one another and because the gene encoding SecB was among the set of 142 genes identified by TraDIS (**figure 2A**), we attempted to construct a Δ*ascA* Δ*secB* double mutant in order to investigate whether AscA and SecB have overlapping functions *in vivo*. Consistent with our TraDIS results, construction of a BW25113 Δ*ascA* Δ*secB* double mutant was only possible when the *ascA* gene was complemented from a high-copy number plasmid. For reasons that are not clear, this strain grew much more poorly than the Δ*secB* single mutant under all conditions tested. However, it was possible to construct a Δ*ascA* Δ*secB* double mutant in *E. coli* MG1655, which is derived from a different lineage of *E. coli* K-12. *E. coli* MG1655 and the Δ*ascA* single mutant grew normally under all conditions tested, and the Δ*secB* mutant displayed a mild cold-sensitive growth defect, consistent with previous studies (25). The Δ*ascA* Δ*secB* double mutant grew normally at 37°C but was extremely cold sensitive for growth (**supplemental figure S4**), indicating that the Δ*ascA* mutation enhanced the phenotype of the Δ*secB* mutant. In addition, overproduction of AscA from a plasmid suppressed the cold sensitive growth defect of the Δ*secB* mutant at 21°C in both BW25113 (**supplemental figure S5**) and MG1655. Finally, the MG1655 Δ*ascA* Δ*secB* double mutant formed filaments of 6-8 cell lengths before ceasing growth when grown at 21°C (**figure 2B**), consistent with a defect in cell envelope biogenesis. Taken together, these results suggested that AscA and SecB have overlapping functions *in vivo*.

**Figure 2.**
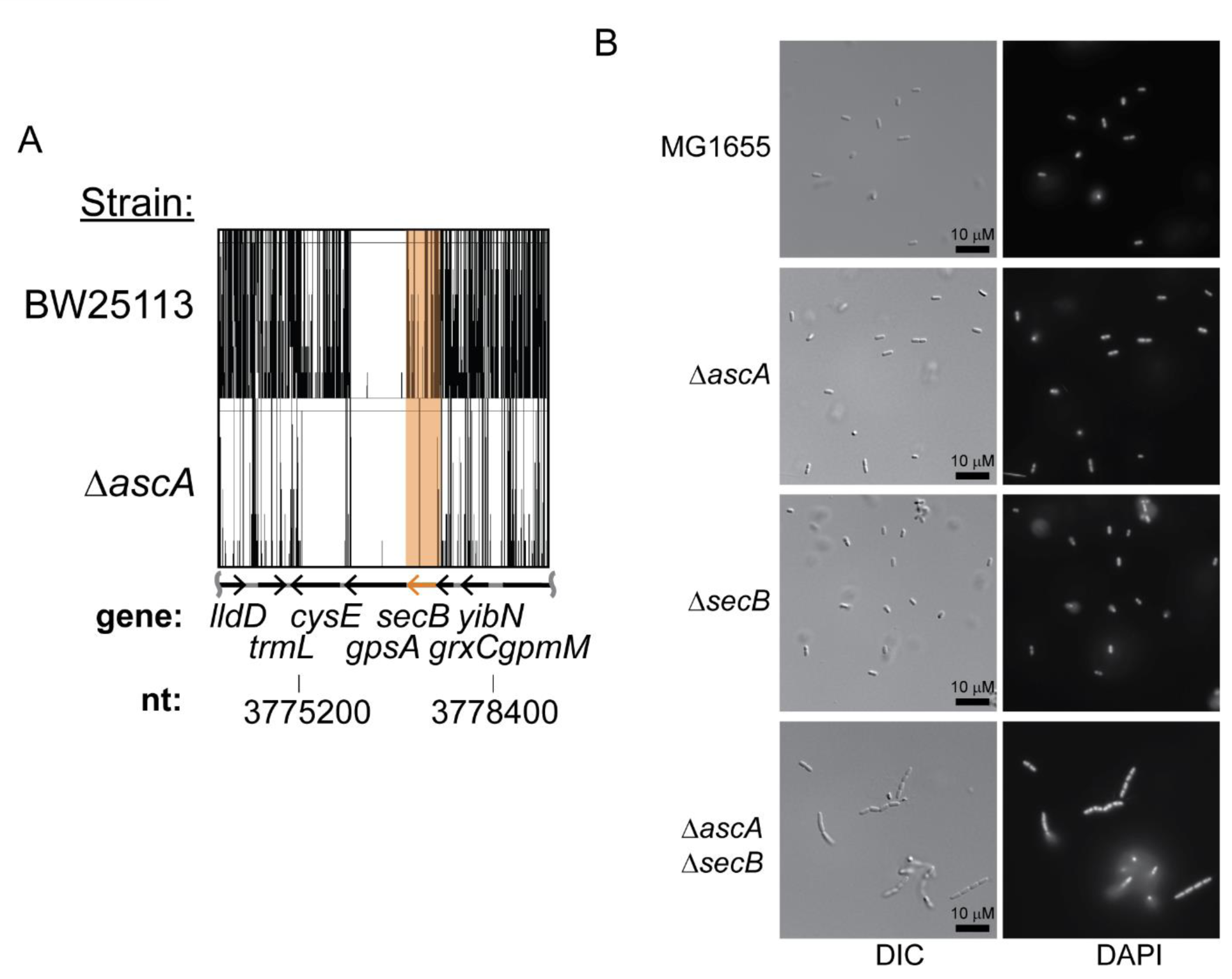
Deletion of the *ascA* gene enhances the cell envelope biogenesis defect of *secB* mutants. (A) A mini-Tn*5*-Kan transposon was hopped into the chromosome of BW25113 Δ*ascA* >500,000 times independently. The transposon insertion sites were determined using Illumina sequencing and compared to insertion sites in a library of transposon insertion mutants in the parent strain (22). Depicted is the region of the chromosome corresponding to nucleotides 3,774,300 to 3,789,300, which contains the *secB* gene (highlighted in orange). Transposon insertion sites in BW25113 (above) and BW25113 Δ*ascA* (below) are indicated by vertical lines. (B) MG1655 (an ancestor of BW25113), MG1655 Δ*ascA*, MG1655 Δ*secB* and MG1655 Δ*secB* Δ*ascA* were diluted into LB and grown at 21°C for 4 hours. Cells were stained with DAPI and imaged using fluorescence microscopy.

### Co-purification of proteins with AscA

To investigate whether AscA interacts with other proteins *in vivo*, we purified a fusion protein between AscA and the small ubiquitin-like modifier from *Saccharomyces cerevisiae* (SUMO), which was tagged at its N-terminus with a Strep(II) affinity tag (Strep-AscA). We then identified the co-purifying polypeptides using mass spectrometry (LC-MS/MS). SDS-PAGE analysis of purified Strep-AscA indicated that it copurified with polypeptides with a range of molecular weights (**supplemental figure S6**). LC-MS/MS analysis of the copurifying polypeptides revealed that full-length AscA copurified with ∼134 different protein species (**figure 3A**), including both cytoplasmic and periplasmic proteins (**supplemental table S2**). However, Strep-AscAΔMBD copurified with only 32 proteins, which was similar to the number of proteins (28) that copurified with a control protein (SUMO from *Saccharomyces cerevisiae*) (**figure 3A**). These results suggested that AscA binds promiscuously to a range of proteins *in vivo* when overexpressed and that the MBD is required for efficient binding to these proteins.

**Figure 3.**
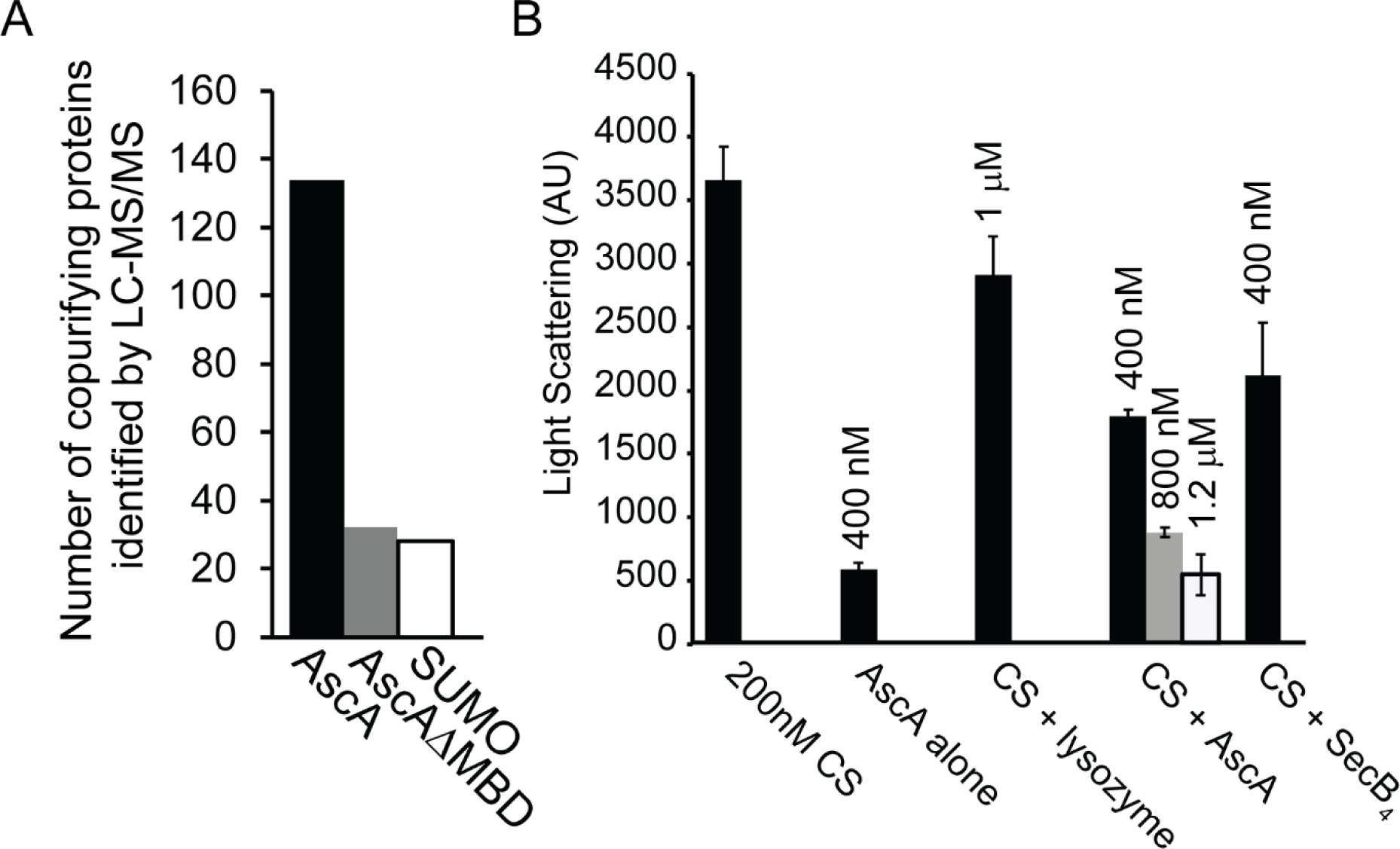
AscA behaves like a molecular chaperone *in vivo* and *in vitro*. (A) A fusion protein between SUMO from *Saccharamyces cerevisiae* and AscA (SUMO-AscA), AscAΔMBD (SUMO-AscAΔMBD) or SUMO alone was purified from lysates of BL21(DE3) by virtue of an N-terminal Strep(II) tag. The co-purifying proteins were then resolved by SDS-PAGE and identified by mass spectrometry (LC-MS/MS). (B) 200 nM porcine citrate synthase (CS) was incubated at 50°C for 30 minutes in the indicated concentration of hen egg lysozyme, AscA or SecB (tetrameric). The amount of aggregation in the sample was then determined using light scattering at 320 nm.

### Effect of AscA on protein aggregation

The promiscuous binding AscA to multiple proteins *in vivo* suggested that it could be a molecular chaperone. To investigate this possibility, we examined the ability of AscA to inhibit the aggregation of citrate synthase (CS) at 50°C using light scattering, a widely used assay for molecular chaperone function (26) (**figure 3B**). The presence of AscA strongly inhibited aggregation of CS in a concentration-dependent manner, suggesting that it is a molecular chaperone (**figure 3B**). SecB also inhibited aggregation of CS, albeit to a lesser extent than AscA. However, hen-egg lysozyme, a thermostable control protein, did not inhibit aggregation of CS.

### Effect of Δ<u>ascA</u> mutation on translocation of MalE-LacZ

To investigate the role of AscA in protein translocation, we examined the effect of a Δ*ascA* mutation on the activity of a reporter protein fusion between MalE and LacZ (MalE-LacZ) (27–29). The MalE portion of the protein targets it for Sec-dependent translocation across the cytoplasmic membrane.

However, translocation results in inactivation of LacZ (β-galactosidase). A Δ*secB* mutation caused an increase in β-galactosidase activity, consistent with previous studies and consistent with the role of SecB in translocation of MalE-LacZ (29) (**figure 4A, black bars**). A Δ*ascA* mutation also caused an increase in β-galactosidase activity, suggesting that defect in AscA inhibit the translocation of MalE-LacZ. Expression of AscA from a plasmid complemented the translocation defect of the Δ*ascA* mutant, and overexpression of AscA in the parent strain reduced the residual β-galactosidase activity below background levels (**figure 4A, grey bars**). However, overexpression of AscA caused an increase in β-galactosidase activity in a Δ*secB* mutant (**figure 4A**), suggesting that AscA actually inhibits translocation in the absence of SecB. Despite apparently enhancing the translocation defect of the MM18 Δ*secB* mutant, overproduction of AscA suppressed the cold-sensitive growth defect.

**Figure 4.**
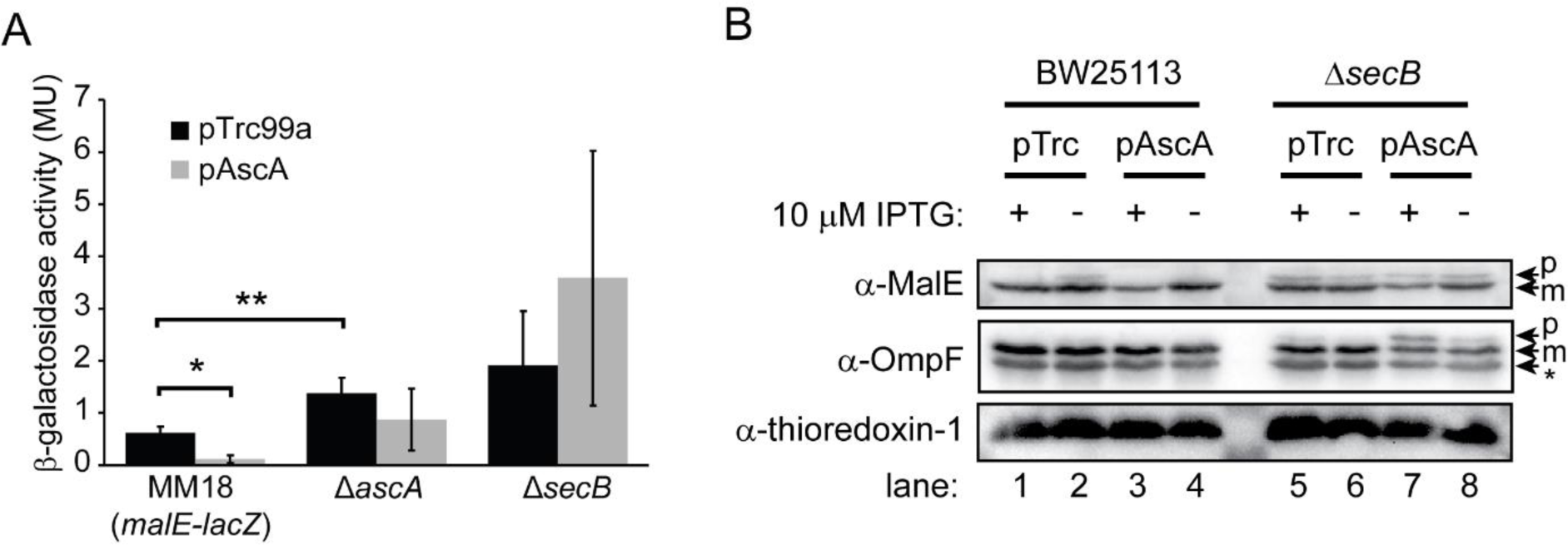
The effect of AscA on Sec-dependent protein translocation *in vivo* is dependent on SecB. (A) *E. coli* MM18 (which produces a MalE-LacZ reporter fusion protein), MM18 Δ*ascA* and MM18 Δ*secB* were transformed with the plasmid vector pTrc99a or pTrc99a containing IPTG-inducible copy of *ascA* (pAscA). Cells grown on LB containing 10μM IPTG, and the defective translocation of MalE-LacZ was determined by measuring the β-galactosidase activities of the strains. (B) BW25113 and BW25113 Δ*secB* containing pTrc99a or pAscA were grown on LB containing 2% maltose. The cell lysates were then resolved by SDS-PAGE and analysed by western blotting using anti-sera against MalE, OmpF and thioredoxin-1. The running positions of precursor-length MalE and OmpF (p), which contains the N-terminal signal sequence, and processed mature-length MalE and OmpF (m), which lack their signal sequences, are indicated.

### Effect of AscA overexpression on Sec substrate proteins

To confirm the effect of AscA on Sec-dependent protein translocation, we examined the effect of AscA overproduction on the translocation of two native model Sec substrates, MalE and OmpF (28). Newly synthesised periplasmic and outer membrane proteins, including MalE and OmpF, contain an N-terminal signal sequence that allows it to be recognised by the Sec machinery (5, 30). Because the signal sequence is removed during translocation, the steady-state levels of precursor proteins (which contain a signal sequence) provides insight into the efficiency of translocation *in vivo*. AscA overproduction did not affect the relative levels of precursor and mature length MalE or OmpF in BW25113 (**figure 4B, lanes 1 - 4**), indicating that it does not cause a translocation defect when expressed on its own. Overproduction did cause a decrease in the steady state levels of MalE in BW25113, consistent with the reduced β-galactosidase activity of strains producing MalE-LacZ, but did not affect the levels of OmpF or thioredoxin-1 (a cytoplasmic control protein). However, overexpression of AscA enhanced the accumulation of precursor-length MalE and OmpF in a Δ*secB* mutant (**figure 4B, lanes 5 - 8**).

### Effect of AscA overexpression of MalE-LacZ toxicity

To investigate the timing of the interaction of AscA with substrate proteins, we examined the ability of AscA to suppress the toxicity of high-level expression of MalE-LacZ. High-level expression of MalE-LacZ is toxic because the LacZ portion of the fusion protein “jams” the essential SecYEG complex (8, 27). We reasoned that AscA could suppress this toxicity if it binds to MalE-LacZ before it engages SecYEG. To this end, we examined the effect of inducing the expression of MalE-LacZ on minimal plates using maltose. Induction of MalE-LacZ in strains containing pTrc99A caused a severe growth defect, consistent with the MalE-LacZ toxicity. However, co-induction of AscA from a high-copy number plasmid relieved this toxicity (**supplemental figure S7**), suggesting that AscA likely binds to its substrate proteins before they engage SecYEG.

## DISCUSSION

Our results suggest that AscA is a ribosome-associated molecular chaperone that assists Sec-dependent protein translocation. When overexpressed, AscA binds promiscuously to a broad range of proteins, and the ribosome-binding activity of AscA suggests that it recognises these proteins while they are still nascent polypeptides. AscA also binds to SecB and facilitates protein translocation *in vivo*. However, overexpression of AscA enhances the translocation defect caused by disruption of the *secB* gene. These results suggest a pathway for AscA-mediated targeting (**figure 5**): (i) AscA recognises nascent substrate proteins as they emerge from the ribosome; (ii) AscA holds proteins in an unfolded conformation in the cytoplasm until (iii) it can target the bound protein to SecB; (iv) SecB then delivers the substrate protein to SecA for translocation through SecYEG across the cytoplasmic membrane.

**Figure 5.**
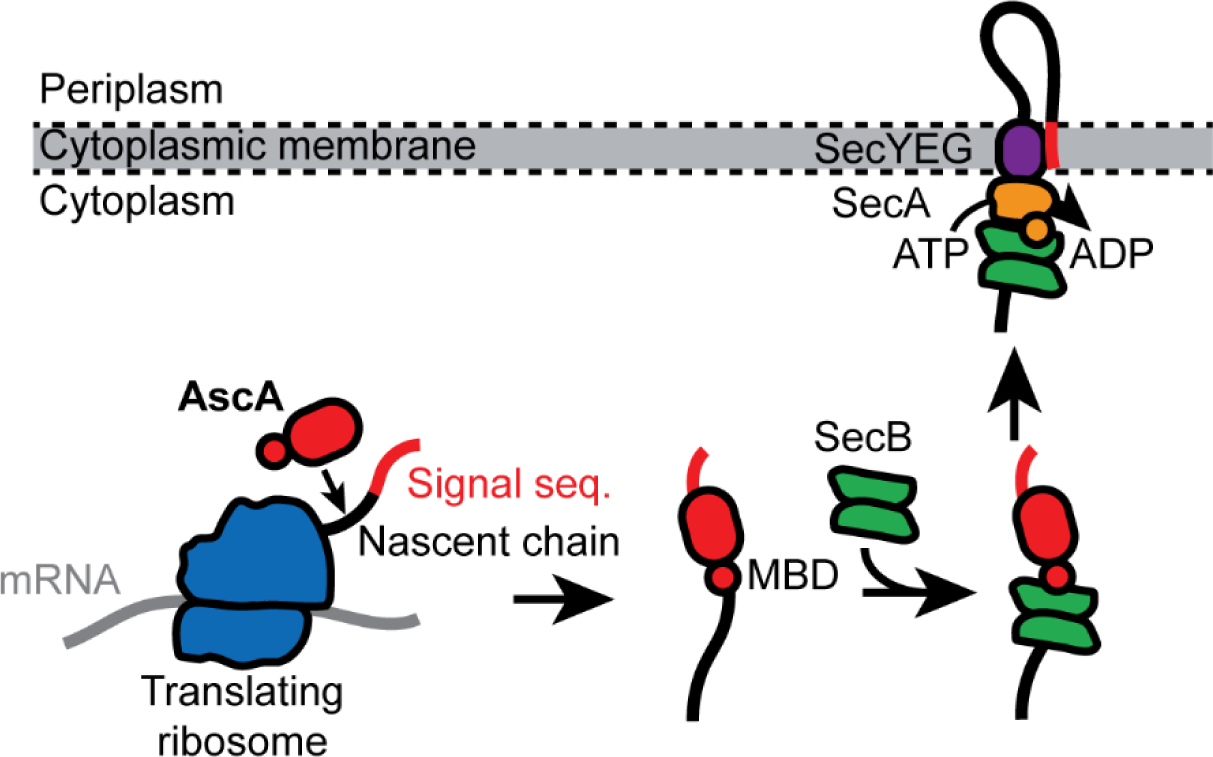
Diagrammatic model of putative role of AscA in Sec-dependent protein translocation. AscA binds to ribosomes, presumably to recognise nascent substrate proteins as they emerge from the ribosome. AscA binds to newly synthesised substrate proteins in the cytoplasm. AscA then targets the substrate protein for Sec-dependent protein translocation by recruiting SecB. SecB then delivers the substrate protein for translocation by binding to SecA.

Expression of AscA appears to be conditional, which could explain why AscA has not been identified in previous genetic screens for Sec components. The conditions required for *ascA* induction are not known, but our results suggest that expression could be regulated by the sRNA RybB, which is regulated by σ^E^, and RhyB, which is regulated by iron availability (24). Our results suggest that AscA binds promiscuously to a range of substrate proteins, and it is possible that AscA recognises all or most nascent SecA substrates. This possibility would allow SecB to bind to nascent substrate proteins even under conditions that limit the availability of free SecA (14). However, the physiological substrates likely depend on the conditions under which AscA is normally expressed.

Overexpression of AscA simultaneously enhances the defect in protein translocation and suppresses the cold sensitive growth in strains defective for SecB, suggesting that the cold sensitivity of Δ*secB* mutants is an indirect effect (25, 31, 32). For example, it is possible that defects in SecB could increase SecYEG jamming at low temperatures, and binding of AscA to these proteins could prevent them from engaging SecYEG while simultaneously enhancing the apparent Sec defect caused by the absence of SecB. Alternatively, the reduction in the steady state levels of MBP upon overexpression of AscA suggests that AscA could be a quality control component that targets excess Sec substrate proteins for degradation. It is possible that AscA is involved in the turnover of Sec components and substrate proteins by Lon protease (32, 33). Finally, our results raise the possibility that the cold sensitive growth defects caused by defect in many Sec components are also an indirect effect of the defect in Sec-dependent protein translocation (31, 34).

The discovery of AscA highlights the importance of molecular chaperones in promoting Sec-dependent protein translocation. In addition to SecB and AscA, which facilitate translocation by retarding the rate of protein folding (9, 35), the ribosome-associated chaperone Trigger Factor prevents nascent outer membrane proteins from engaging the Sec machinery cotranslationally (36). Other housekeeping chaperones, such as the DnaKJ and GroEL/ES systems, can also facilitate Sec-dependent protein translocation when over expressed (37, 38). However, the physiological importance of these systems in Sec-dependent protein translocation is unknown. It has been suggested that some periplasmic chaperones assist protein translocation by inhibiting retrograde translocation (39). Our results raise the possibility the size and importance of the network of molecular chaperones that support Sec machinery is much greater than is currently known.

The high sequence conservation of SecA-like MBDs suggests that their function is to facilitate interactions with the Sec machinery and with ribosomes (18). Both SecA and AscA MBD also binds to both SecB and ribosomes, and these interactions, except for the interaction of SecA with the ribosome, are dependent on the MBD (15, 17). Although AscA is not universally conserved, 92 of the 156 representative species used for a recent analysis of SecA produce an MBD-containing protein besides SecA (40). Many of these species produce multiple MBD-containing proteins besides SecA. For example, *E. coli* produces a third MBD-containing protein, YchJ (18), and *Salmonella typhimurium* produces five MBD-containing proteins, including SecA and AscA. Furthermore, in 17 of these species, the corresponding SecA protein does not contain an MBD. Besides SecA and AscA, the functions of these proteins are unknown, but bioinformatic analysis suggests at least 18 different functions associated with MBD-containing proteins (41–44). We speculate that SecA-like MBDs are part of an evolutionary toolkit that, when fused to a protein, can bring new functions to the bacterial Sec machinery. If true, MBD-containing proteins could belong to a network of accessory components that assist, modify and/or enhance Sec-dependent protein translocation.

## METHODS

### Chemicals and media

All chemicals were purchased from Sigma-Aldrich (St. Louis, MO, USA) unless otherwise indicated. Rabbit anti-AscA antiserum was produced using purified AscA by Eurogentec (Liège, Belgium). Rabbit anti-OmpF antiserum was a kind gift from T. Silhavy. Rabbit anti-thioredoxin-1 antiserum was purchased from Sigma-Aldrich.

Horseradish peroxidase (HRP)-labelled anti-rabbit antibody was purchased from GE Healthcare. Strains were typically grown in lysogeny broth (LB). Where indicated, strains were grown in M9 minimal medium containing the indicated carbon source (45). Maltose was used at a final concentration of 1% or 0.2%, as indicated, and glycerol was used at a final concentration of 0.5%. Kanamycin and ampicillin were used when required at a final concentration of 30 μg/ml and 200 μg/ml, respectively. When used, 100 μl of a 50 mg/ml solution of 5-bromo-4-chloro-3-indolyl-β-D-galactopyranoside (x-gal) was spread directly onto plates. IPTG and chloroamphenicol were added to the growth media at the indicated concentrations.

### Strains and plasmids

Strains and plasmids were constructed using standard methods (45, 46) and are listed **supplemental table S3**. Plasmids pDH585, pDH963, pCS070, pCS071 and pCS163 were constructed by amplifying the DNA encoding AscA, AscAΔMBD or SecB by PCR and ligating into plasmid pCA528 or pCA597 using the *BsaI* and *BamHI* restriction sites (47). Plasmid pAscA was constructed by amplifying the DNA encoding AscA containing *PciI* and *BamHI* sticky ends and ligating into plasmid pTrc99a (Stratagene) between the *NcoI* and *BamHI* sites. Strain MM18 was a kind gift from J. Beckwith. Single-gene replacement mutations Δ*ascA*::Kan and Δ*secB*::Kan were obtained from the Keio collection(23) and confirmed by PCR.

### Protein purification

Unfused full-length AscA, AscAΔMBD and SecB were purified as described (14, 18). BL21(DE3) containing plasmid pCS070 or pCS071 was grown to mid-log phase in LB containing kanamycin at 37°C, and expression of His_3_-SUMO-AscA (or His_3_-SUMO-AscAΔMBD) was induced using 1 mM isopropyl-thio-galactoside (IPTG; Bioline) overnight at 18°C. Cells were lysed by cell disruption, and lysates were cycled over a 5 ml His-Trap column (GE Healthcare) at 4°C overnight. The column was washed using 25 ml of high-salt wash buffer (20 mM potassium HEPES pH 7.5, 500 mM potassium acetate, 10 mM magnesium acetate, 50 mM imidazole) followed by 25 ml low-salt wash buffer (20 mM potassium HEPES pH 7.5, 25 mM potassium acetate, 10 mM magnesium acetate, 50 mM imidazole). The bound protein was eluted from the column using elution buffer (20 mM potassium HEPES pH 7.5, 25 mM potassium acetate, 10 mM magnesium acetate, 500 mM imidazole). The eluted protein was dialysed by anion-exchange chromatography using a 1ml ResourceQ column (GE Healthcare) equilibrated with 20 mM potassium HEPES pH 7.5, 25 mM potassium acetate, 10 mM magnesium acetate and raising the potassium acetate concentration to 500 mM using a linear gradient over 25 column volumes. The fusion protein was cut with purified His_6_-Ulp1 protease, passaged over a His-Trap column to remove the His-tagged SUMO moiety and concentrated using a VivaSpin20 concentrator with a 10 kDa molecular weight cut-off. Finally, the purified, concentrated protein was cleaned up using using a Superdex S-200 column equilibrated with 20 mM potassium HEPES pH 7.5, 25 mM potassium acetate, 10 mM magnesium acetate.

### Microscale thermophoresis

Purified SecB was randomly labelled using an NT-647-NHS labelling kit (Nanotemper Technologies). 160 nM NT-647-labelled SecB was incubated with the indicated concentrations of unlabelled AscA or AscAΔMBD in buffer (20 mM potassium HEPES [pH 7.5], 100 mM KOAc, 10 mM MgOAc, 0.05% Tween). The thermophoretic properties were of the labelled SecB were then determined with a Monolith NT.115 (Nanotemper Technologies) using MST Premium Coated Capillaries at 100% MST power. Binding constants were determined by fitting the data using the Nanotemper software.

### Ribosome cosedimentation

Cosedimentation of AscA with ribosomes was assayed as described previously (14). Purified AscA was incubated with ribosomes at the indicated concentration in 10 mM HEPES potassium salt, pH 7.5, 25 mM potassium acetate, 10 mM magnesium acetate for >10 minutes. The reaction mixture was then layered on top of a 30% sucrose cushion made with the same buffer and centrifuged at >200,000 x g for 90 minutes. The concentration of ribosomes in the pellet fractions were normalised using the absorbance at 260 nm. Samples were resolved by SDS-PAGE and the amount of AscA in the pellet was determined by western blotting (46).

### Identification of AscA-copurifying proteins

BL21(DE3) containing plasmid pDH963, pCS163 or pCA597 were in LB containing kanamycin grown at 37°C. Expression of Strep-SUMO-AscA, Strep-SUMO-AscAΔMBD and Strep-SUMO was induced at OD_600_ 1.0 with 1 mM IPTG for 2 hours at 30°C. Cells were resuspended in buffer containing 10 mM potassium HEPES (pH 7.4), 100 mM potassium acetate, 10 mM magnesium acetate, 1 mM phenylmethylsulfonyluoride (PMSF) and 10 mg bovine Dnase I and lysed by sonication. The cell debris was removed from the lysate by centrifugation (16000 x *g* for 5 minutes at 4°C), and 1 ml of the clarified lysate was incubated with 50 μl Strep(II)-Tactin Sepharose resin (IBA) for 5 minutes at 4°C. The bound resin was then washed three times with 1 ml 10 mM potassium HEPES (pH 7.4), 100 mM potassium acetate, 10 mM magnesium acetate, three times with 1 ml 10 mM potassium HEPES (pH 7.4), 1000 mM potassium acetate, 10 mM magnesium acetate and one time with 100 mM Tris (pH 8.0) to remove the excess salt. The resin was then dried using a SpeedVac, resuspended in 50 μl SDS-PAGE sample buffer and boiled for 5 minutes. The eluted protein was resolved by SDS-PAGE and Coomassie staining. The co-purifying proteins in the sections of each lane corresponding to molecular weights of greater than ∼80 kDa, ∼60-80 kDa and less than ∼60 kDa were identified using LC/MS-MS.

### Citrate synthase aggregation assay

200 nM porcine citrate synthase (Sigma-Aldrich) was incubated incubated in 10mM potassium HEPES (pH 7.40, 100 mM potassium acetate, 10 mM magnesium acetate in the presence of the indicated concentration of AscA, SecB or hen egg lysozyme (Sigma-Aldrich) at 50°C. After 30 minutes, samples were returned to room temperature, and side scattering of light at 320 nm was determined using a fluorometer.

### TraDIS

TraDIS experiments were conducted as described previously (21, 22). A library of ∼500,000 independent mini-Tn*5* transposon mutants was created by transforming BW25113 Δ*ascA* with an EZ-Tn*5* <Kan-2> Tnp transposome kit (Epicentre) by electroporation. Kanamycin-resistant transformants were pooled by flooding the transformation plates with LB. After extracting the genomic DNA from the pooled colonies using a RTP Bacteria DNA Mini Kit (Stratech, Ely, UK), the DNA concentration was determined by Qubit (Invitrogen) and sheared by sonication. The sheared genomic DNA was prepared for sequencing using a NEBNext Ultra DNA Library Prep Kit for Illumina (New England Biolabs) with an additional PCR step to amplify for transposon containing fragments. The PCR products were purified using the Agencourt AMPure XP system by Beckman Coulter. The products were sequenced using Illumina Miseq V3 (150 cycle) cartridges on an Illumina MiSeq sequencer. The locations of the sequences were mapped to the *E. coli* reference genome CP_009273.1 (*E. coli* K-12 BW25113). Files containing the locations of the mapped sequences can be found at doi: 10.6084/m9.figshare.12676739.

### Microscopy

Cells were grown overnight in LB at 37°C, subcultured into LB, and grown at 21°C for 4 hours to OD_600_ 0.4. Cells were fixed by pelleting and resuspending in PBS containing 2.5% paraformaldehyde. The cells were stained with DAPI and immobilised on a poly-L-lysine-coated glass slide. Cells were imaged using a Zeiss Axio Observer.Z1/7 inverted microscope with a Plan Apochromat 100X Oil DIC M27 objective (numerical aperture 1.4). DIC images were produced using a TL Halogen light source, and epifluoscence images were produced using an Albireo LED lightsource at 405 nm using a 90HE DAPI/GFP/Cy3/Cy5 reflector (maximum emission = 465 nm; maximum excitation = 353 nm). Images were captured using an Axiocam 503m detector and subsequently processed using Zeiss ZEN 2.3 software and ImageJ 1.51j8 (48).

### β-galactosidase assays

Because the β-galactosidase activities of cells grown in culture were variable, strains were streaked onto the indicated plates containing x-gal (which was included to screen against the spontaneous loss of the *malE-lacZ* gene) and IPTG and grown overnight at 37°C. Samples were prepared by scraping cells from the plate using a wire loop and resuspending in Z-buffer (45). Otherwise, the β-galactosidase activities of strains producing MalE-LacZ were determined as described (45).

### Western blotting

To determine the steady-state levels of MalE, OmpF and thioredoxin-1, cells were streaked onto LB plates containing 1% maltose to induce the *mal* regulon and grown at 37°C. After overnight growth, cells were scraped from the plate using a wire loop and resuspended in LB. The samples were then normalised to OD_600_ 1.0 by spectrophotometry, pelleted by centrifugation and resuspended in 100 μl SDS-PAGE sample buffer. The samples were resolved by SDS-PAGE and analysed by western blotting using antisera specific to the indicated protein (46).

## ACKNOWLEDGEMENTS.

We thank J. Cole, D. Grainger, T. Knowles, W. Allen, I. Collinson, J. Bryant, I. Cadby and members of the T101 lab for insightful advice and discussion. TCS, MW and MTM were funded by the Biotechnology and Biological Sciences Research Council (BBSRC) Midlands Integrated Integrative Biosciences Training Partnership (MIBTP). DH and MJ were funded by BBSRC grant BB/L019434/1. We thank MRC-CLIMB for cloud computing access. We thank A. di Maio and the Birmingham Advanced Light Microscopy Facility for assistance with microscopy. We thank the Biosciences Functional Genomics Facility and the Advanced Mass Spectrometry and Proteomics Facility at the University of Birmingham.

---

**Supplemental figure S1.**
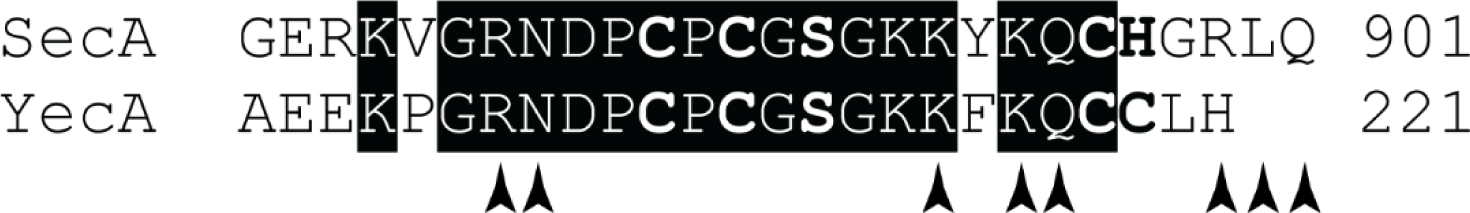
Sequence comparison of the SecA and YecA MBDs. Sequence alignment of the C-terminal MBDs of SecA (above) and YecA (below) from *E. coli* MG1655. The amino acid number of the final amino amino acid is indicated. Amino acids involved in coordinating the bound metal are bolded. Identical amino acids are highlighted in black. Amino acids involved in binding to SecB in the x-ray crystal structure from *Haemophilus influenzae* (1OZB; (20)) are indicated with a carrot below.

**Supplemental figure S2.**
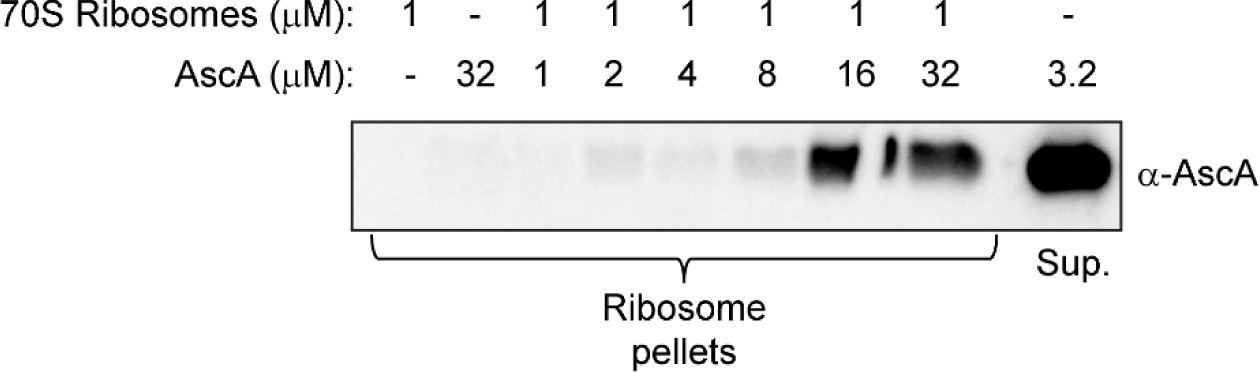
Effect of AscA concentration on cosedimentation with the ribosome. 1 μM vacant 70S ribosomes were incubated with the indicated concentration of AscA, and the binding reaction was layered on top of a 30% sucrose cushion. Ribosomes were then pelleted through the sucrose cushion by ultracentrifugation, the ribosomal pellets were resuspended buffer and the concentrations of the ribosomes in the resulting were normalised using absorbance at 260nm. The samples were then resolved by SDS-PAGE and analysed by western blotting using antiserum against AscA. Purified AscA at a concentration with a ratio of ∼3.2:1 compared to ribosomes was included as a loading control.

**Supplemental figure S3.**
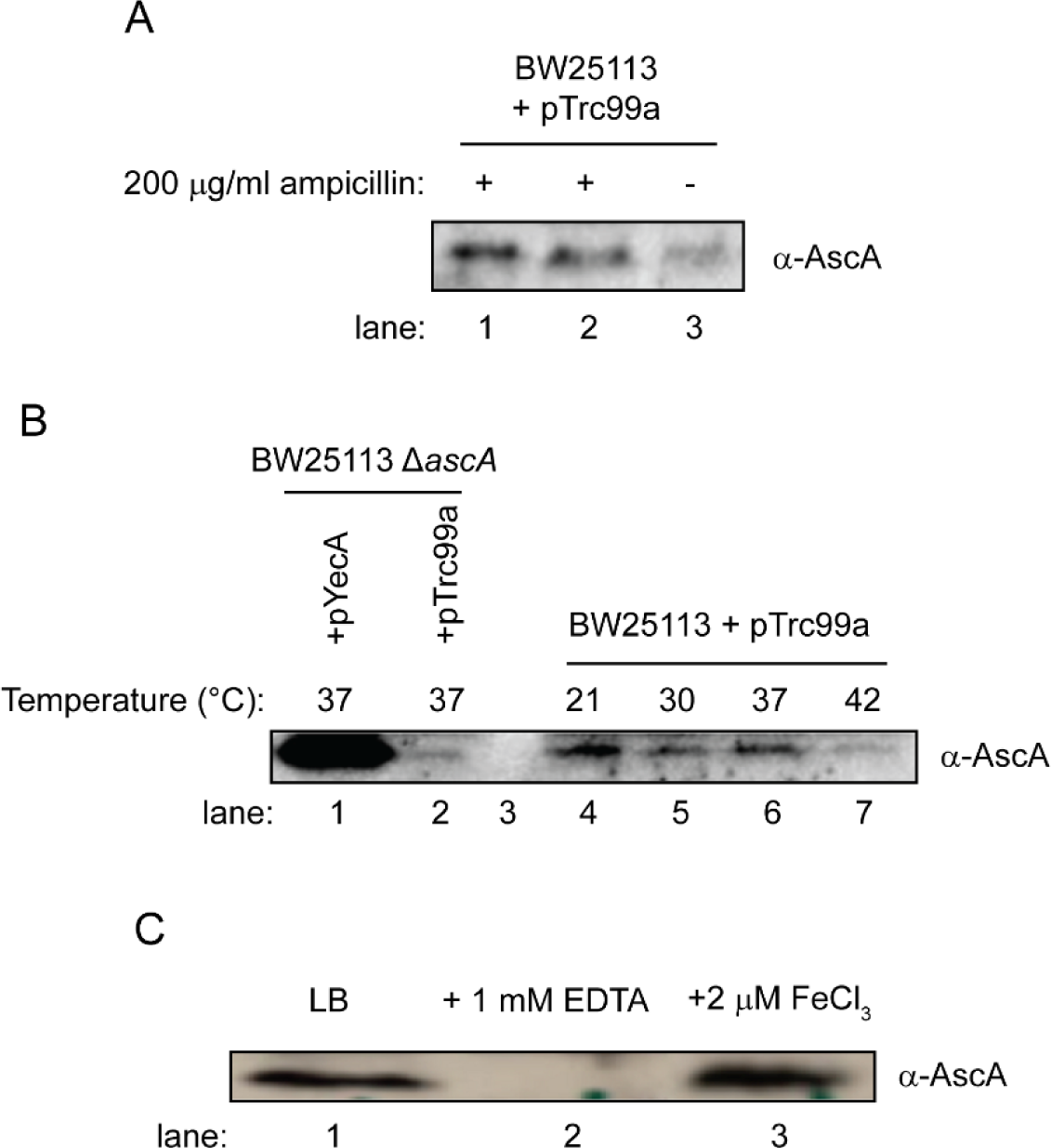
Effect of growth conditions on expression of AscA. The steady state levels of AscA in BW25113 were determined by western blotting. 10 μl of cell lysate normalised to OD_600_ 10 in 1X Laemmli buffer were resolved by SDS-PAGE and analysed by western blotting against AscA. Each experiment was repeated at least three independent times to ensure that differences between samples were not the result of differences in loading. (A) BW25113 containing plasmid pTrc99a (an empty vector containing an ampicillin resistance marker) was grown in the presence (+) or absence (-) of 200 μg/ml ampicillin. (B) BW25113 containing pTrc99a was grown in LB with ampicillin at 21°C (lane 4), 30°C (lane 5), 37°C (lane 6) or 42°C (lane 7). BW25113 Δ*ascA* containing pAscA (lane 1) or pTrc99a (lane 2) were included as controls. (C) BW25113 containing pTrc99a was grown in LB (lane 1), LB containing 1 mM EDTA (lane 2) or LB containing 2 μM FeCl_3_.

**Supplemental figure S4.**
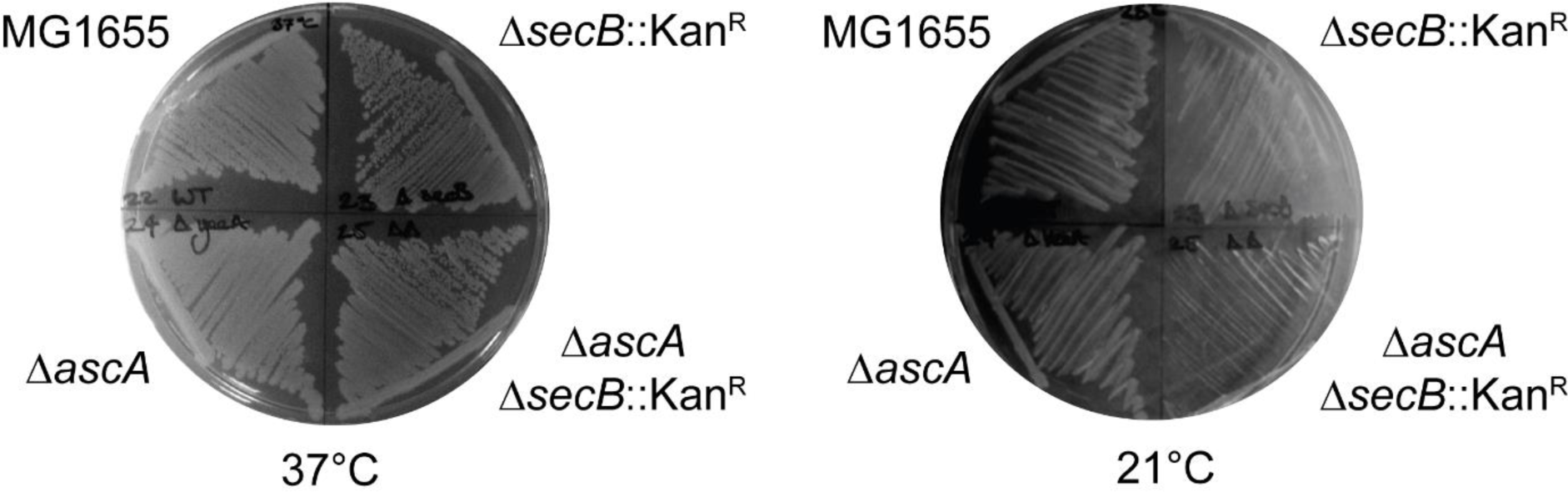
Deletion of the *ascA* gene enhances the cold sensitivity of a Δ*secB* mutant. MG1655, DRH959 (Δ*secB*::Kan^R^), DRH974 (Δ*ascA*), DRH975 (Δ*ascA* Δ*secB*::Kan^R^) were streaked onto LB plates and incubated overnight at 37°C or 48 hours at 21°C.

**Supplemental figure S5.**
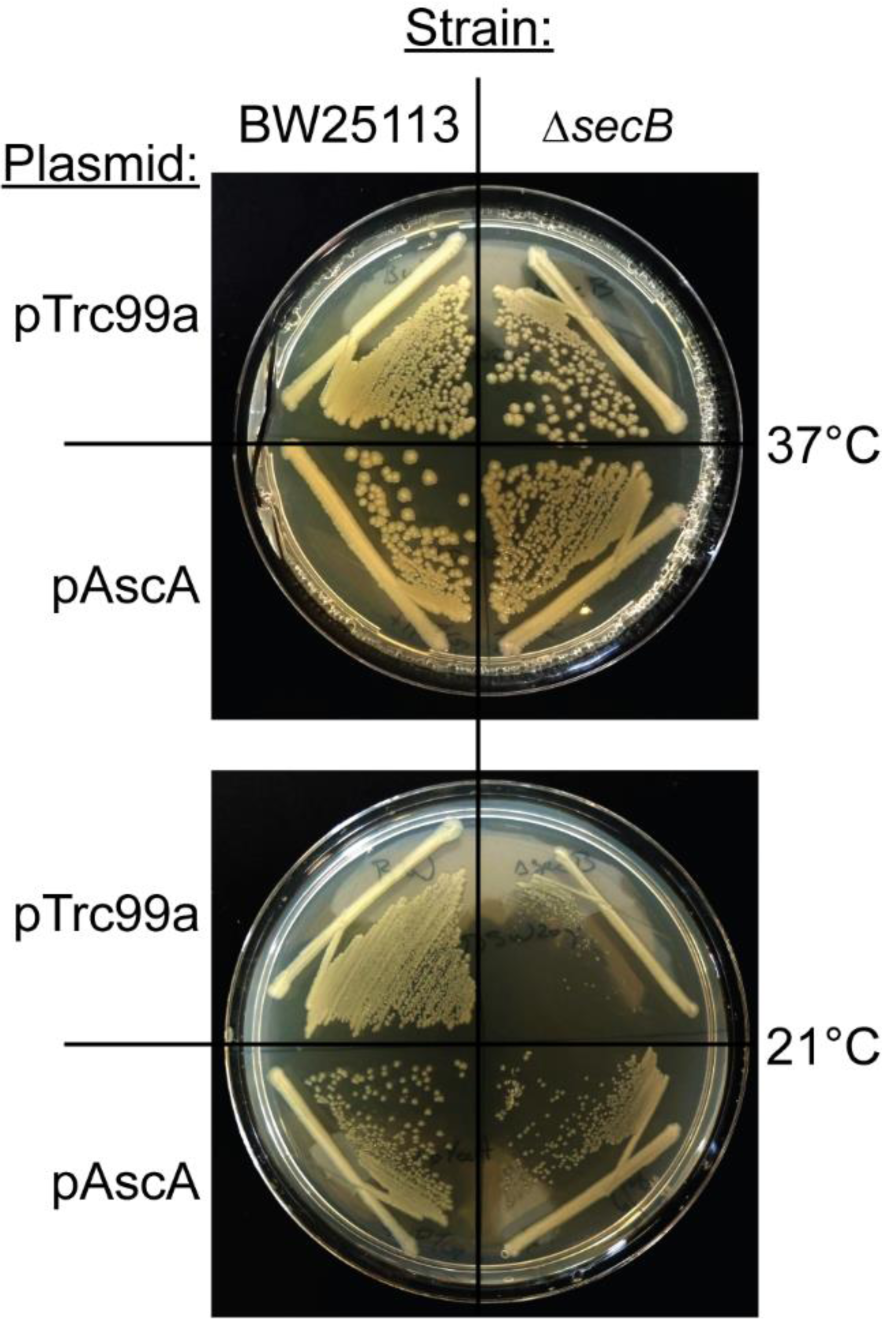
AscA overexpression suppresses the cold-sensitive growth defect caused by a Δ*secB* mutation. BW25113 or BW25113 Δ*secB* containing pTrc99a or pAscA were streaked on LB plates containing 10 μM IPTG and grown overnight at 37°C or for 48 hours at 21°C.

**Supplemental figure S6.**
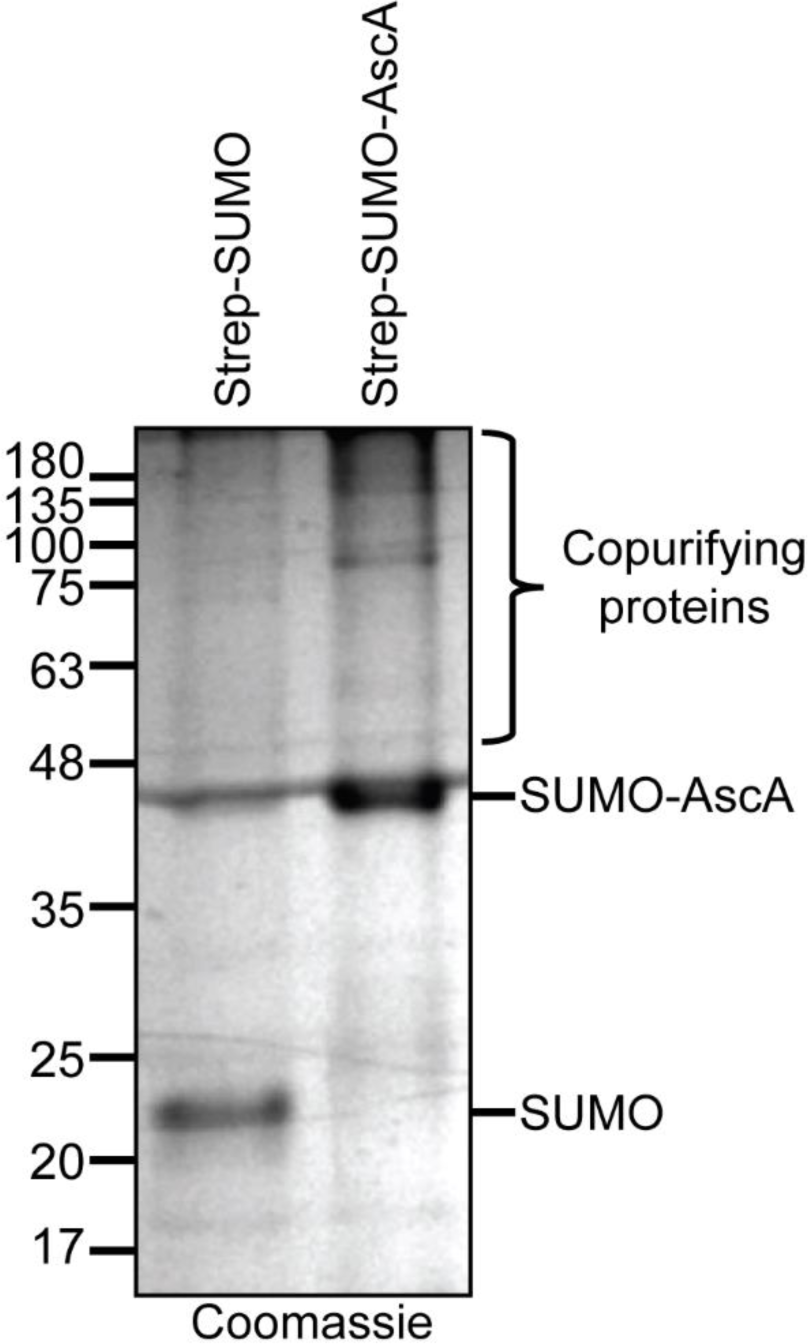
SUMO-AscA copurifies with proteins with a range of molecular weights. Strep(II)_3_-tagged SUMO from *Saccharamyces cerevisiae* (Strep-SUMO) or a SUMO-AscA fusion protein (Strep-SUMO-AscA) was produced in BL21(DE3), purified by affinity chromatography against the Strep tag and analysed by SDS-PAGE and Coomassie staining. A large number of co-purifying proteins with a range of molecular weights, which copurified with Strep-SUMO-AscA but not Strep-SUMO, were detectable by Coomassie staining.

**Supplemental figure S7.**
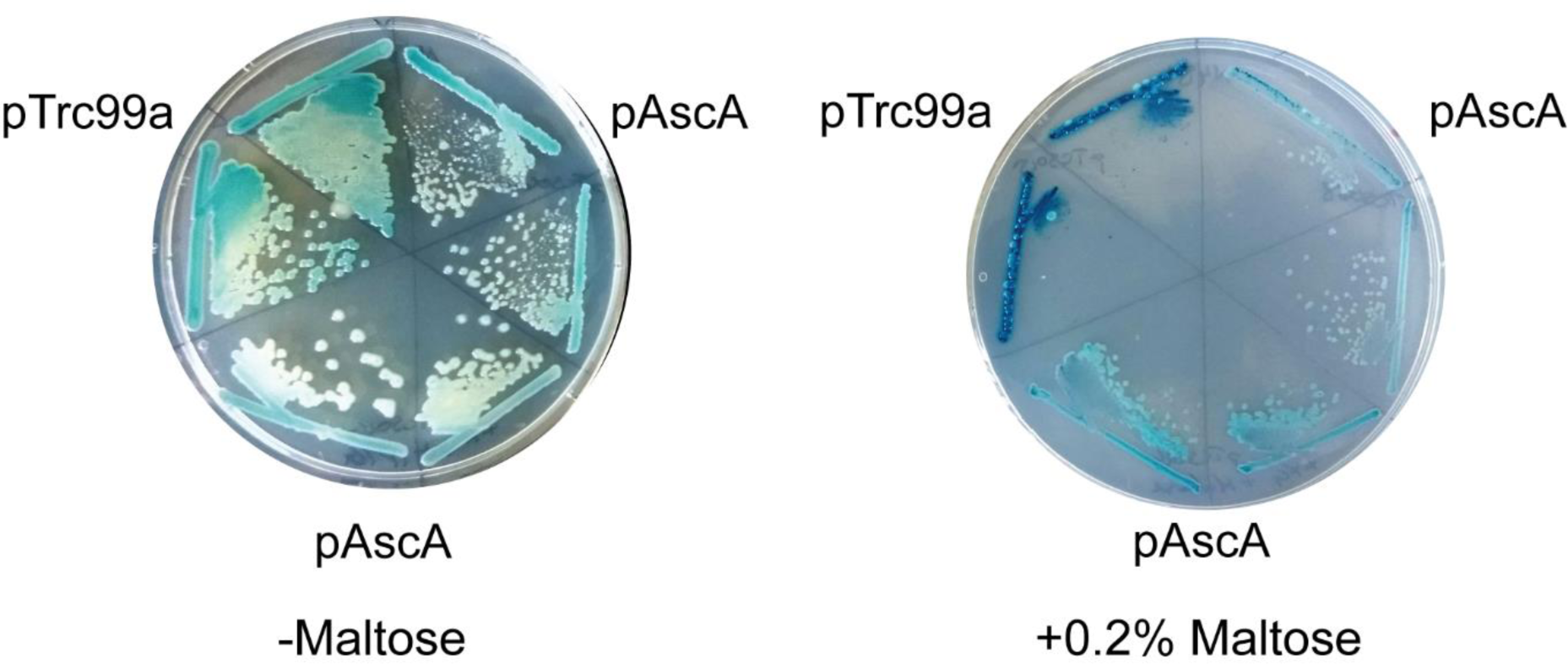
Expression of AscA suppresses the lethality of MalE-LacZ induction. MM18 transformed with pTrc99a or pAscA was streaked onto M63 minimal plates containing 0.5% glycerol, 25 μM IPTG, 200 μg/ml ampicillin and x-gal in the absence (left) or presence (right) of 0.2% maltose to induce high-level expression of MalE-LacZ, which is toxic.

**Supplemental table S1.**
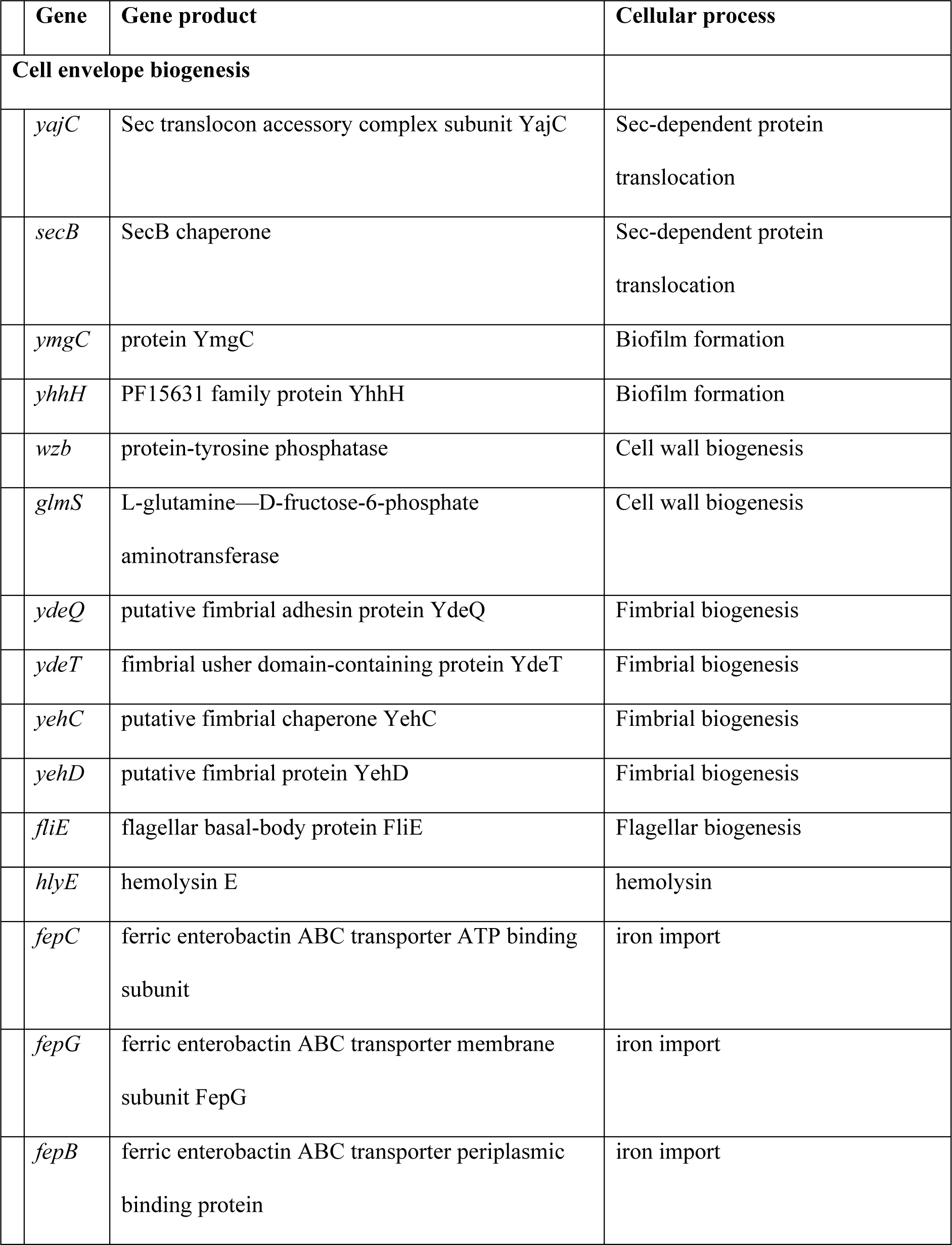
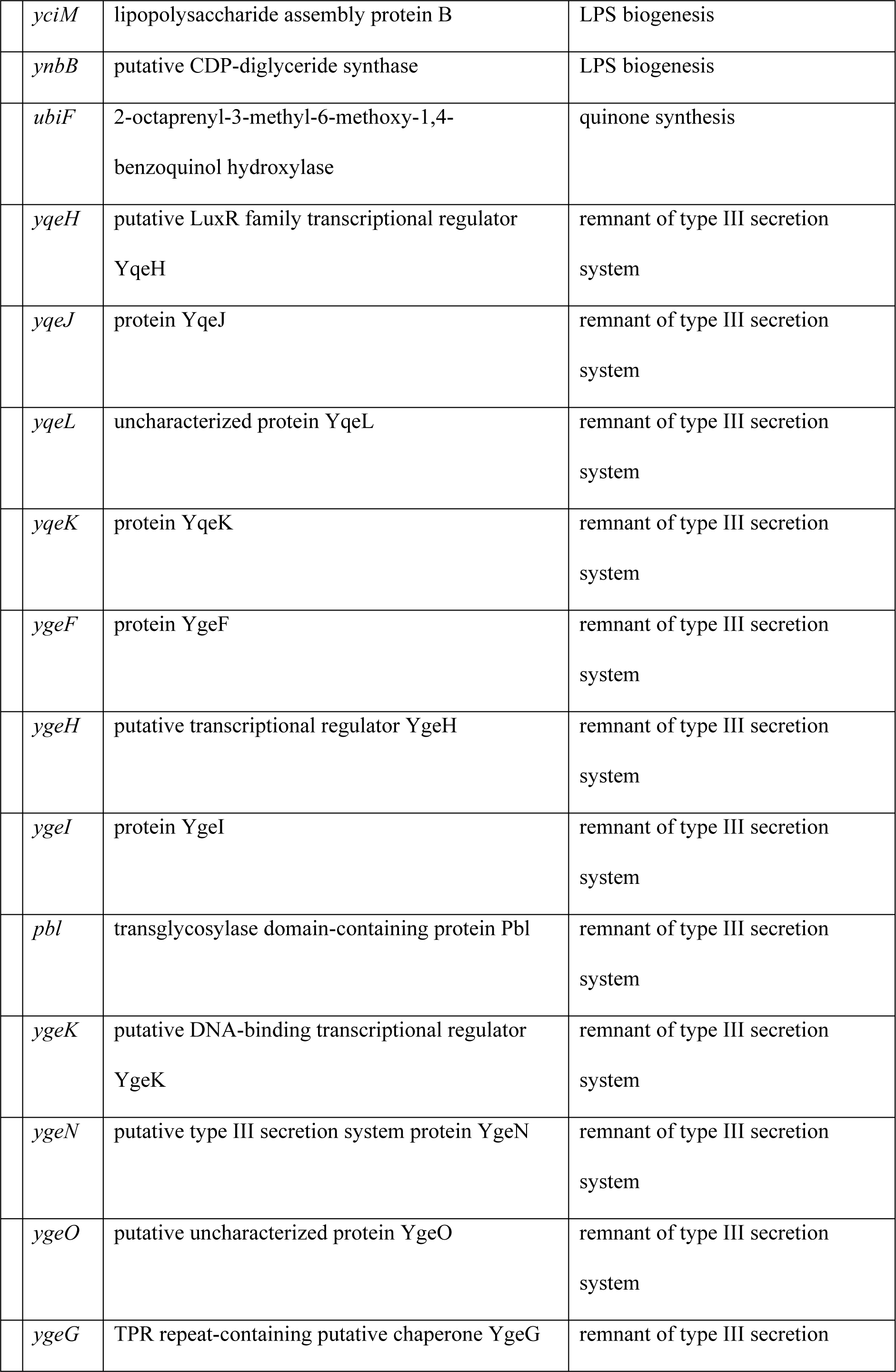
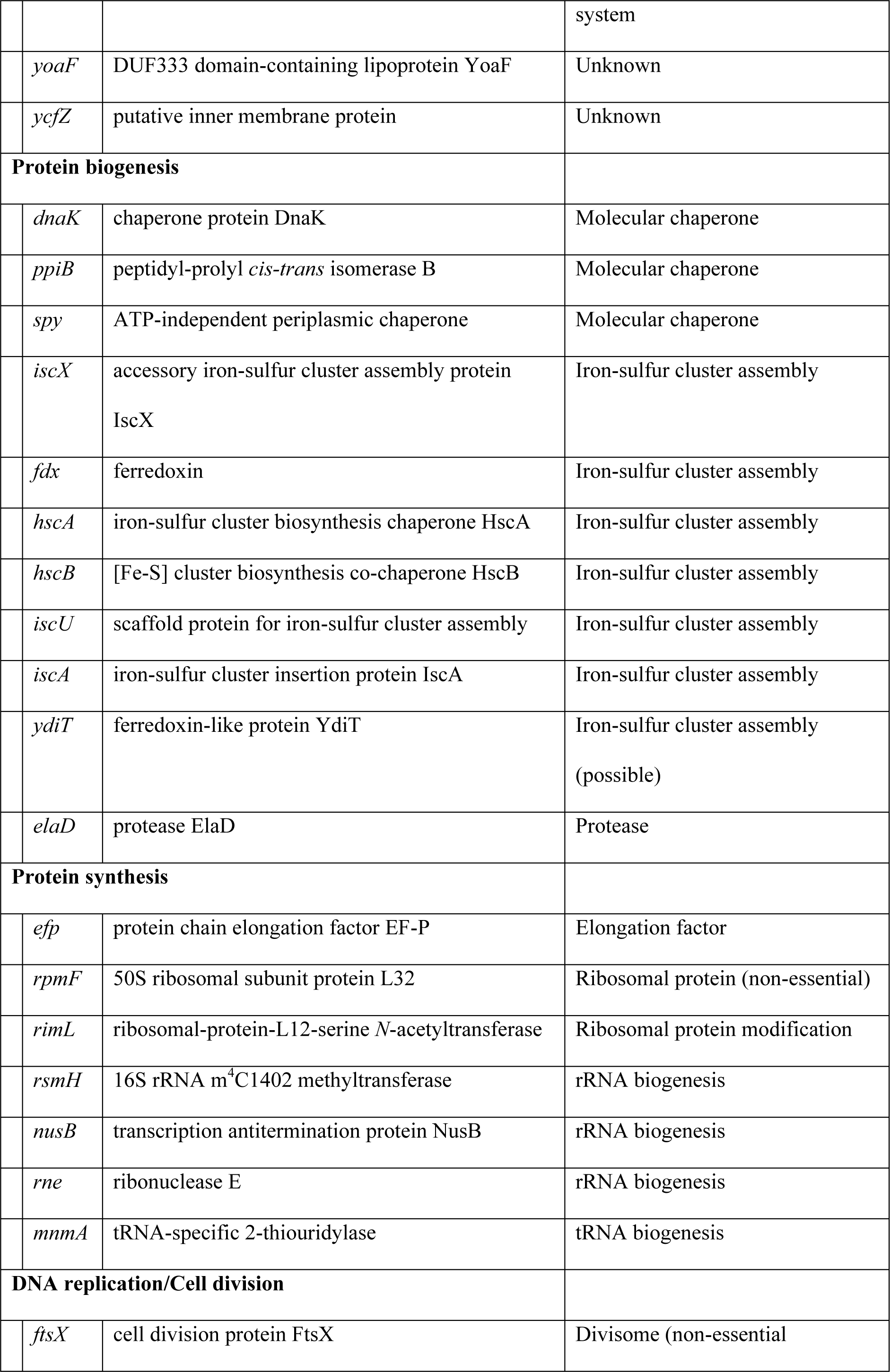
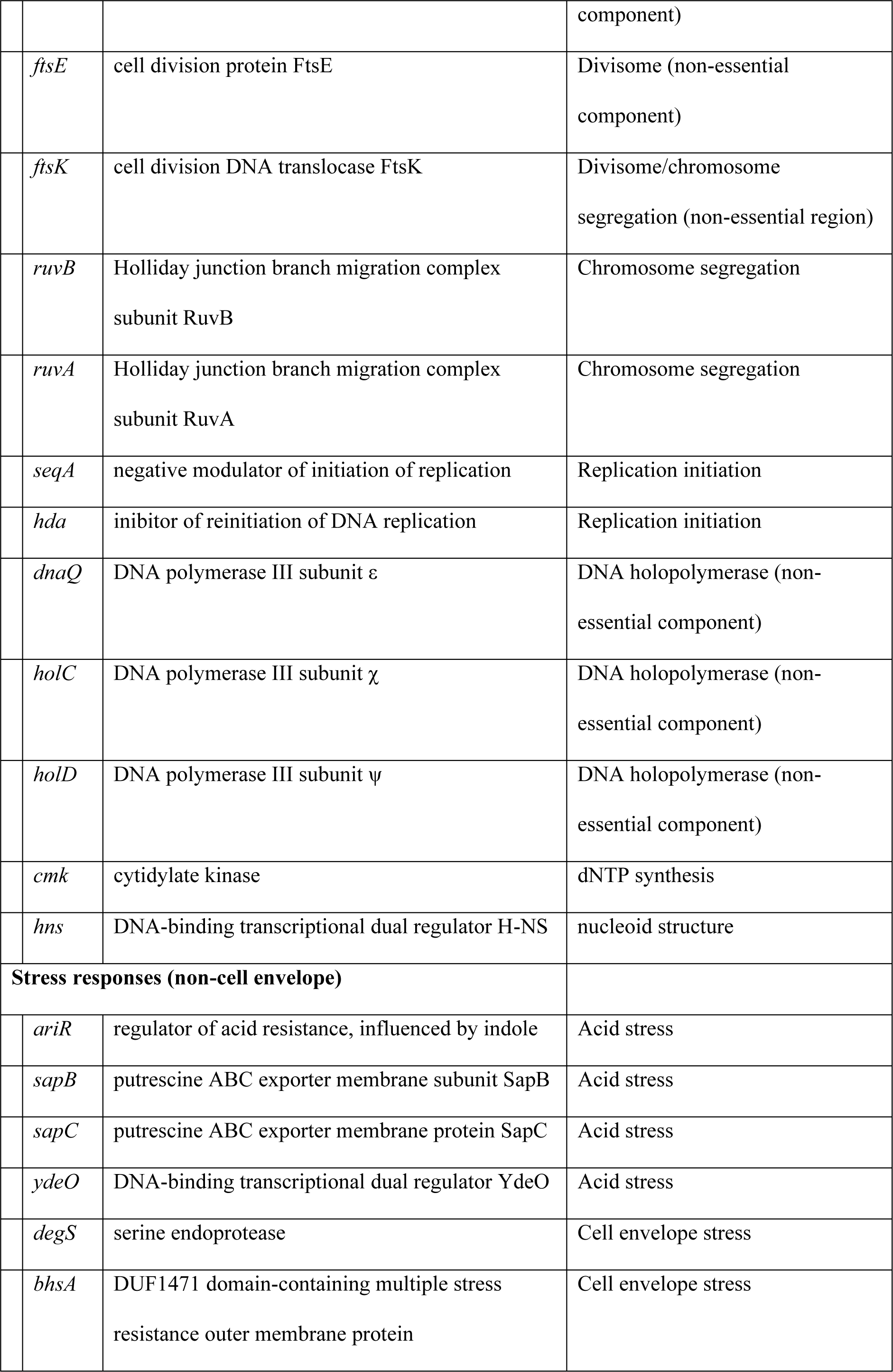
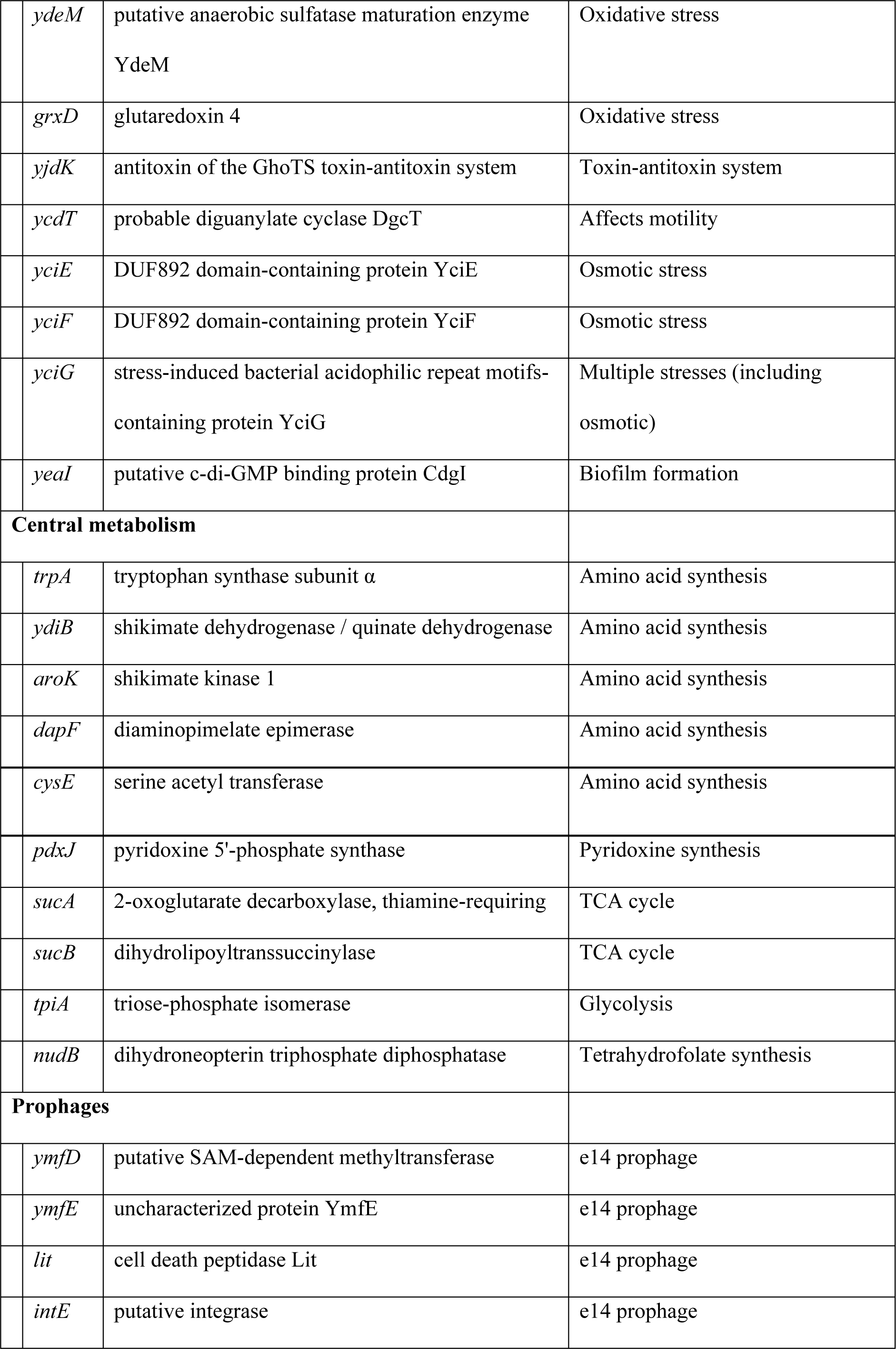
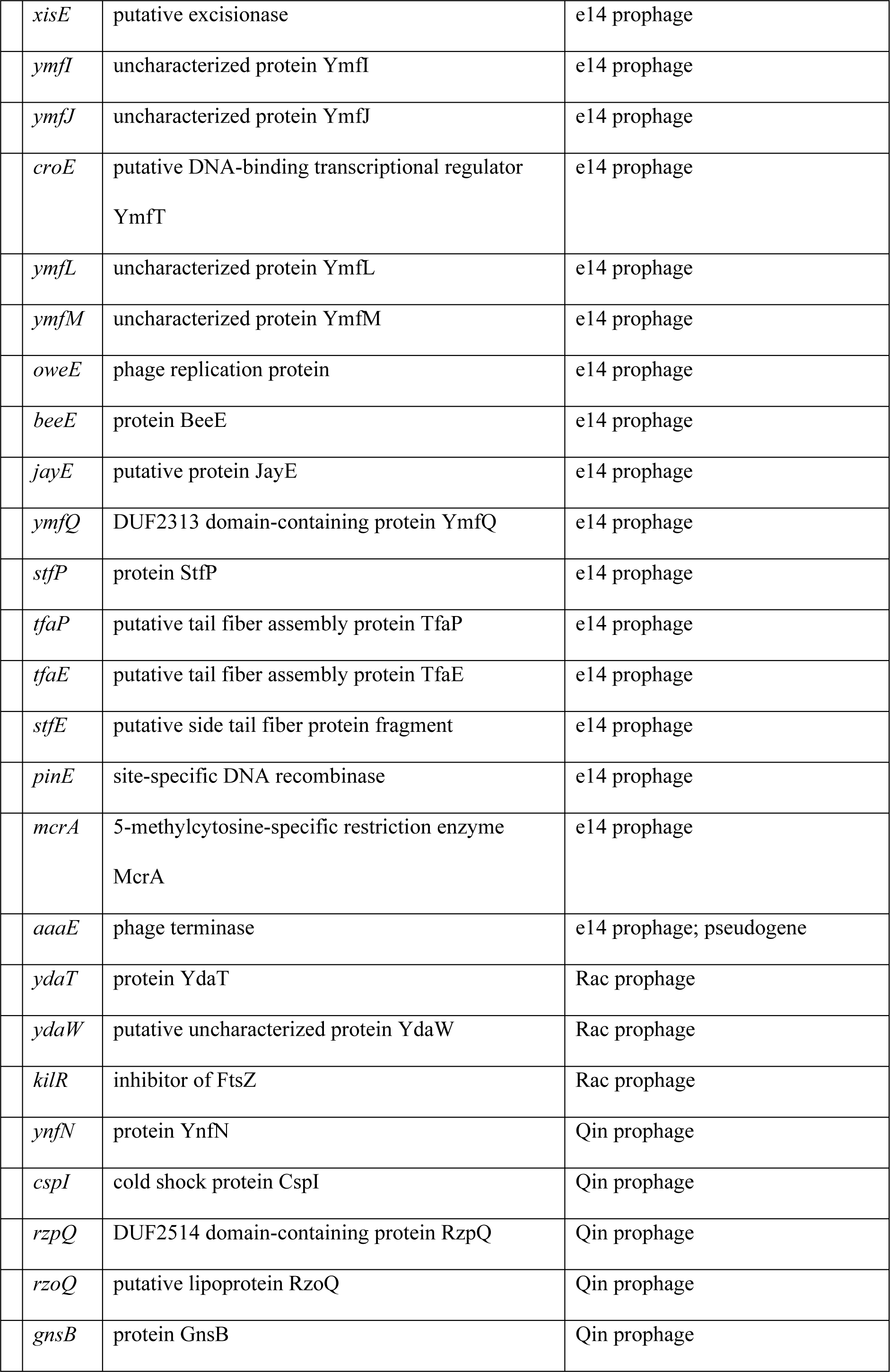
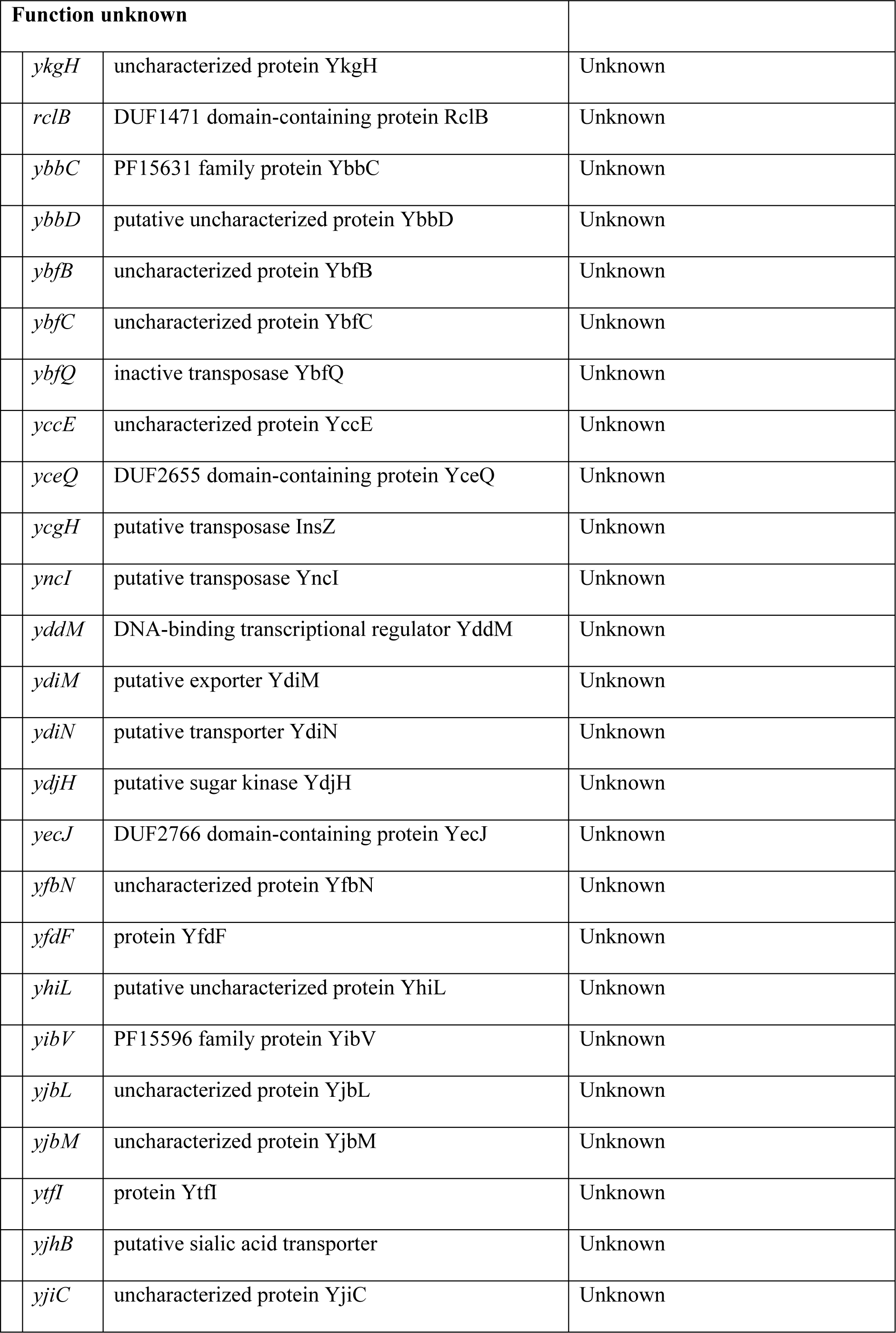
Genes unambiguously identified by TraDIS as essential for viability in BW25113 Δ*yecA*.

**Supplemental table S2.**
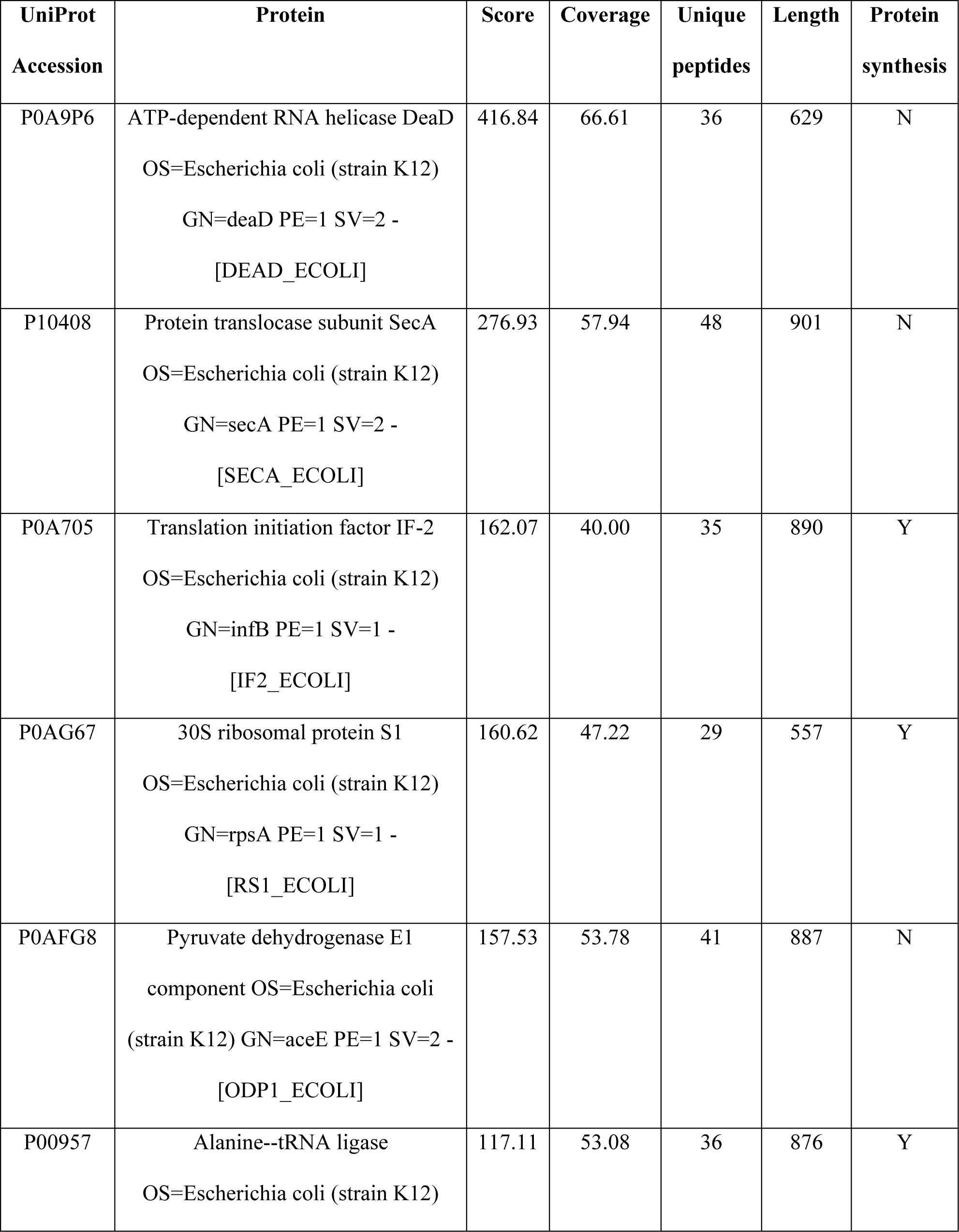
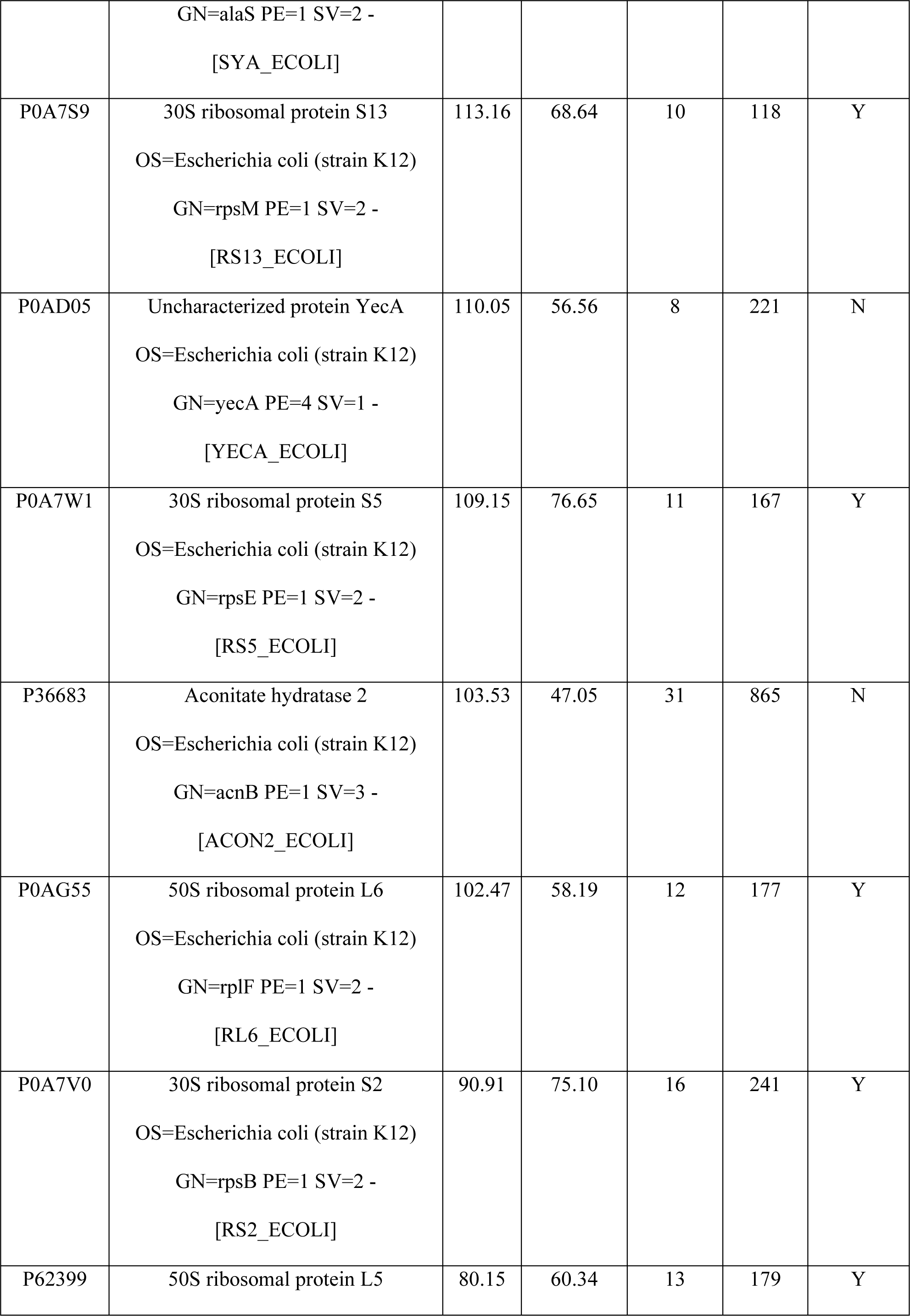
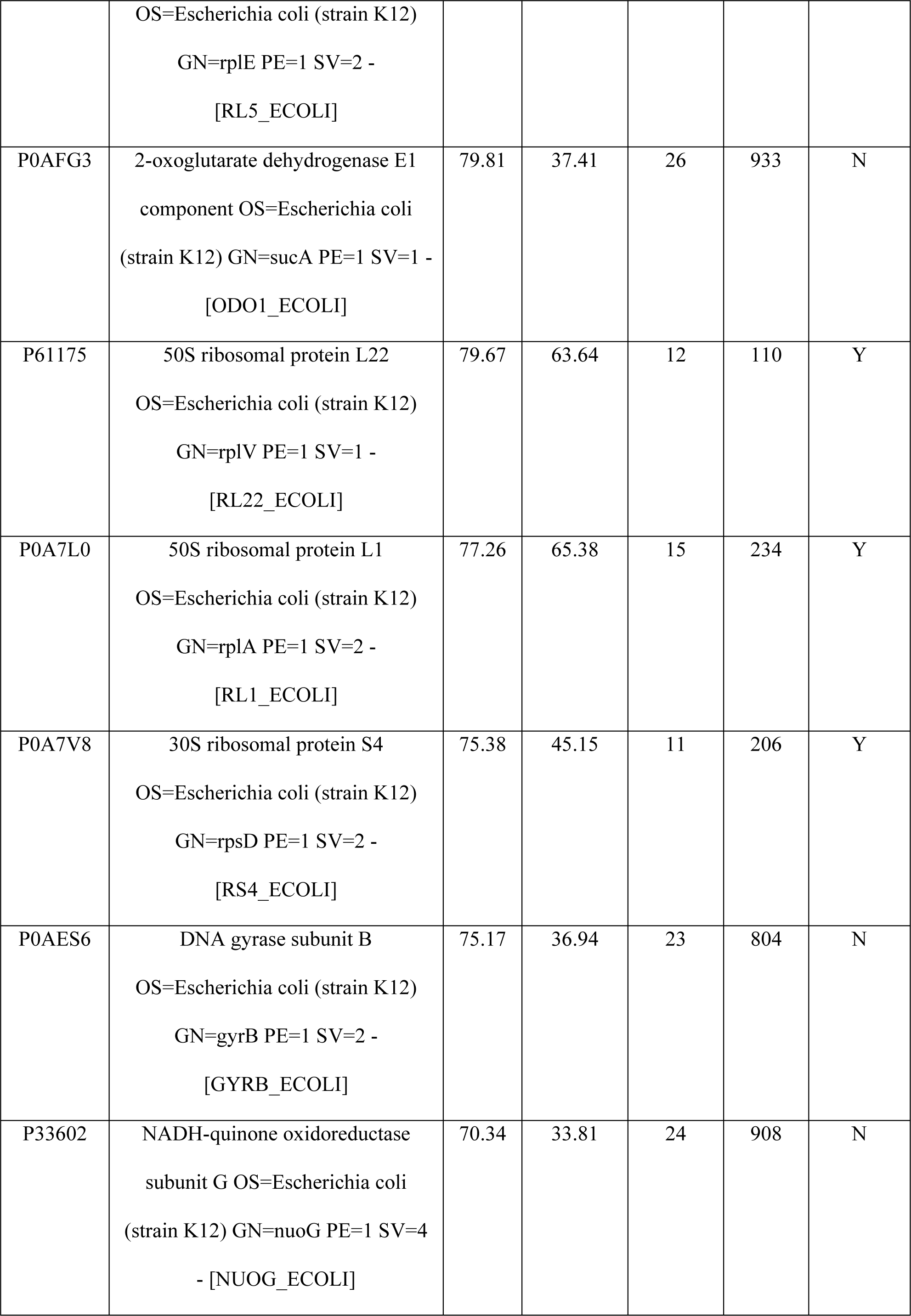
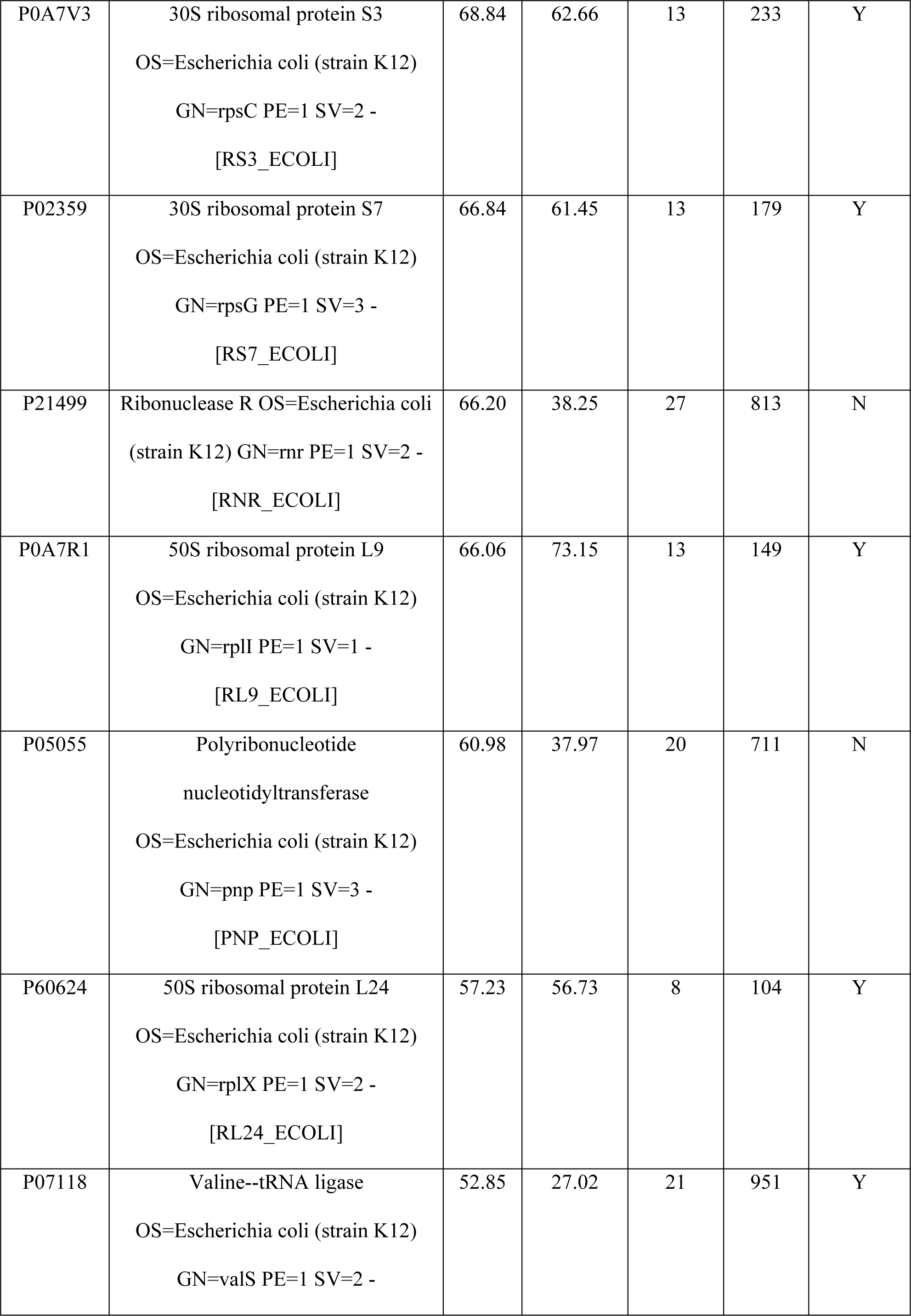
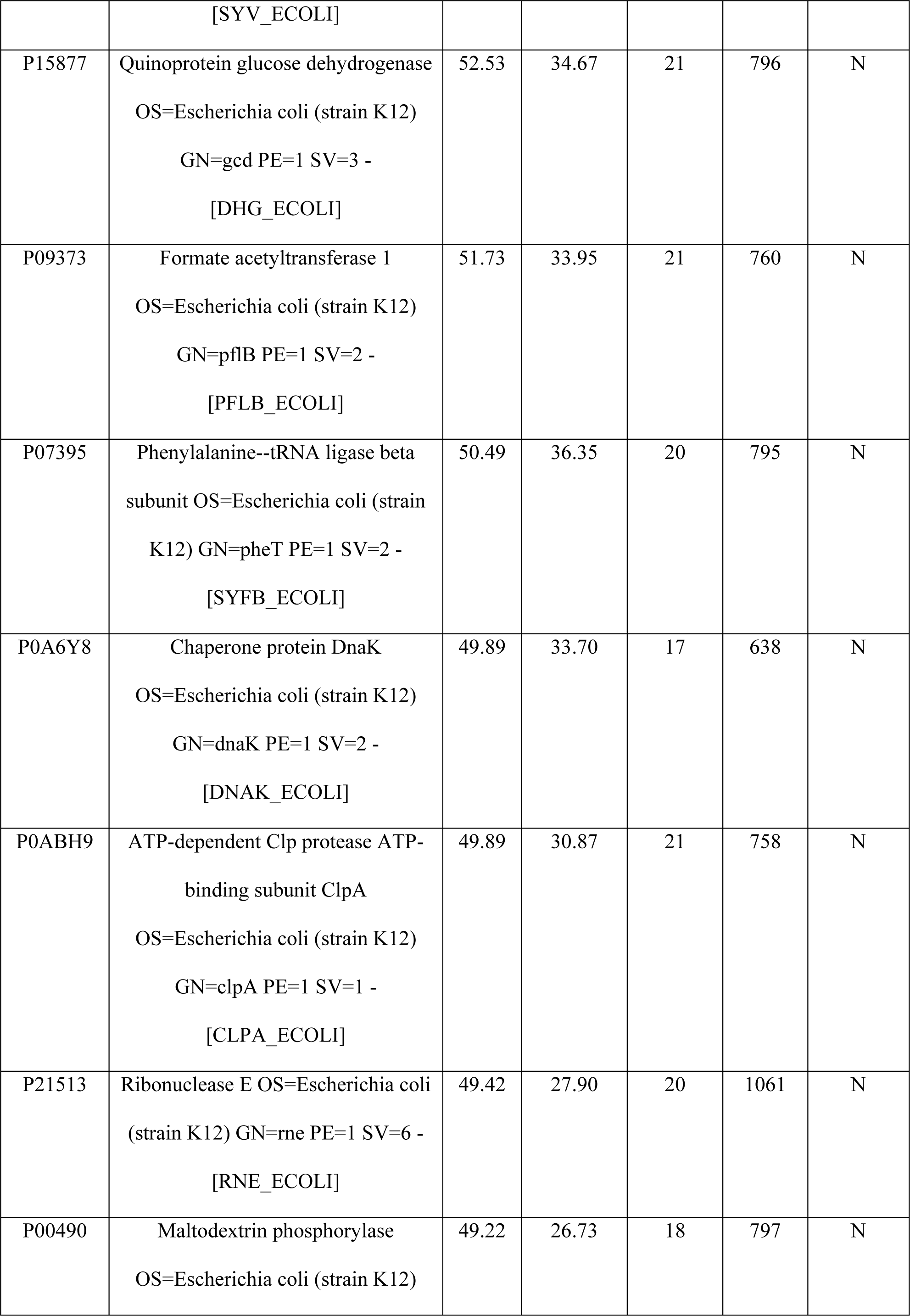
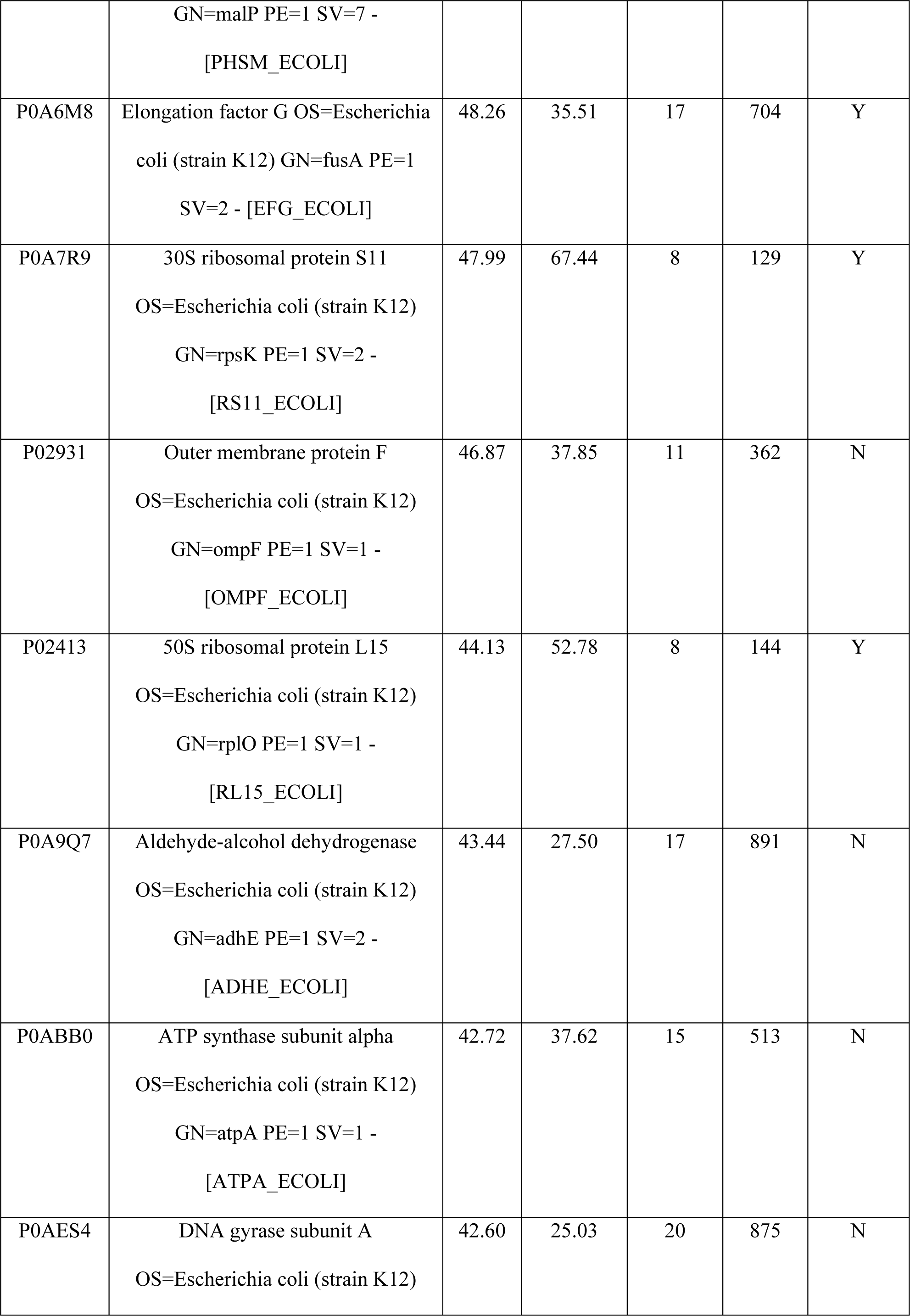
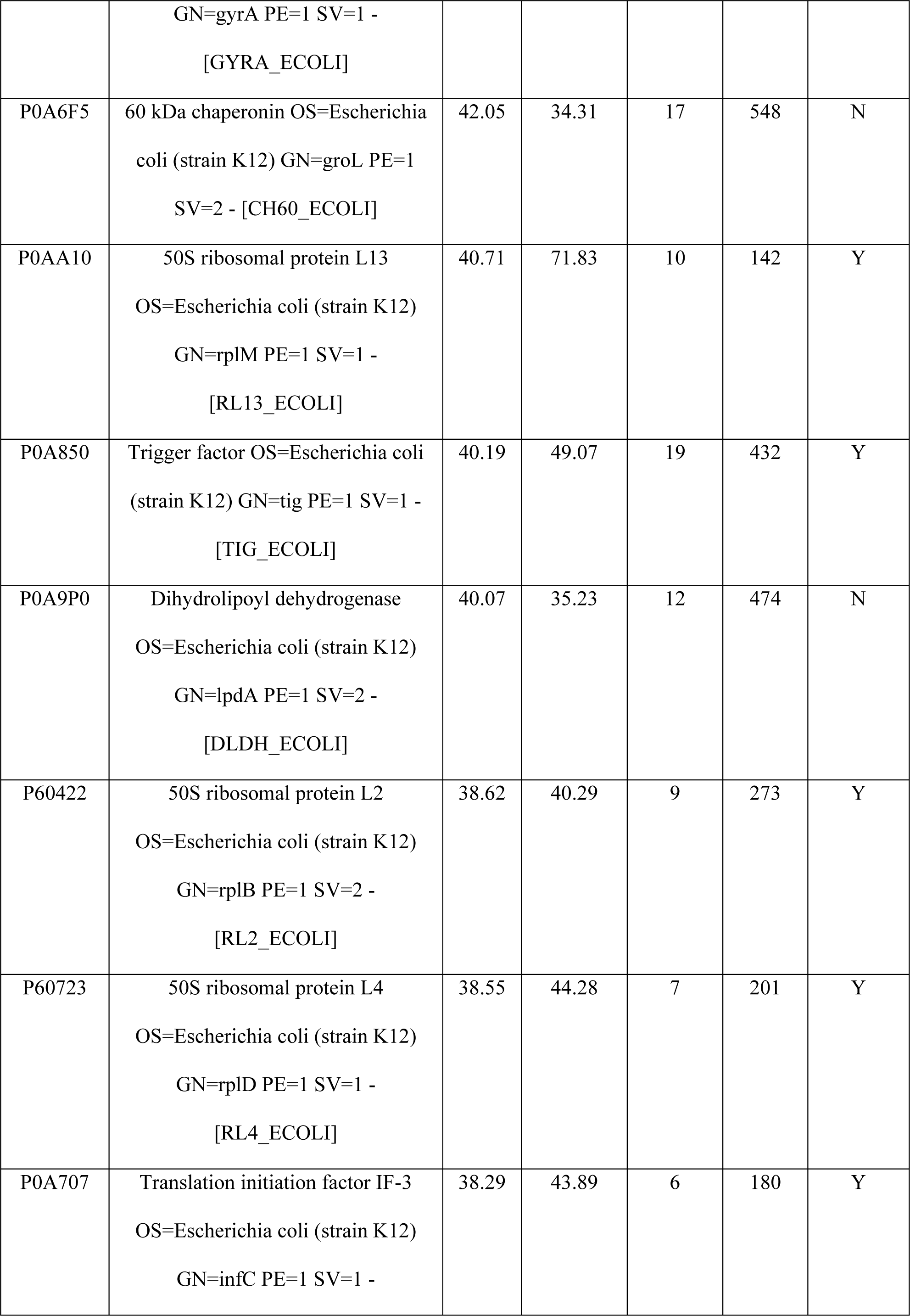
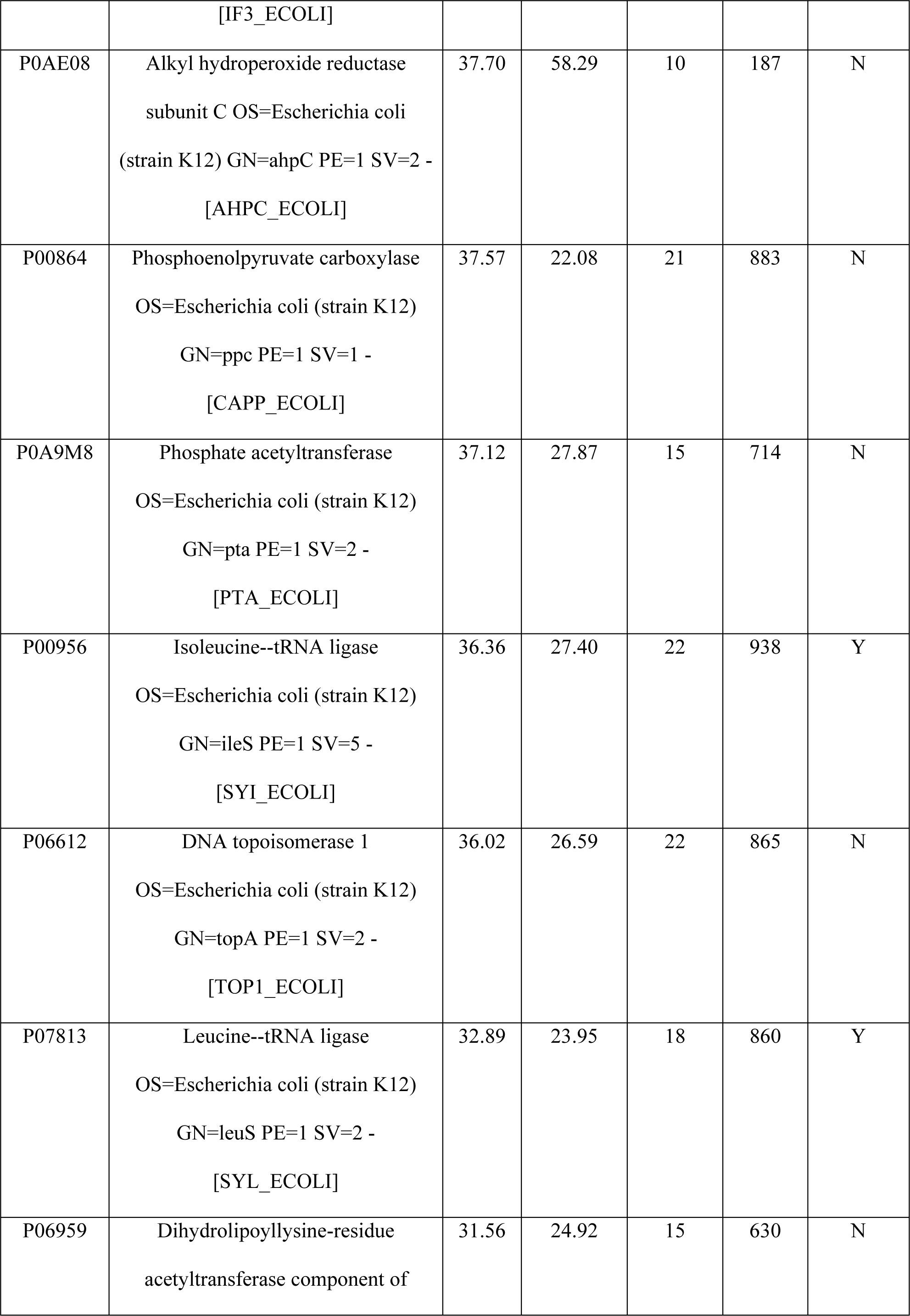
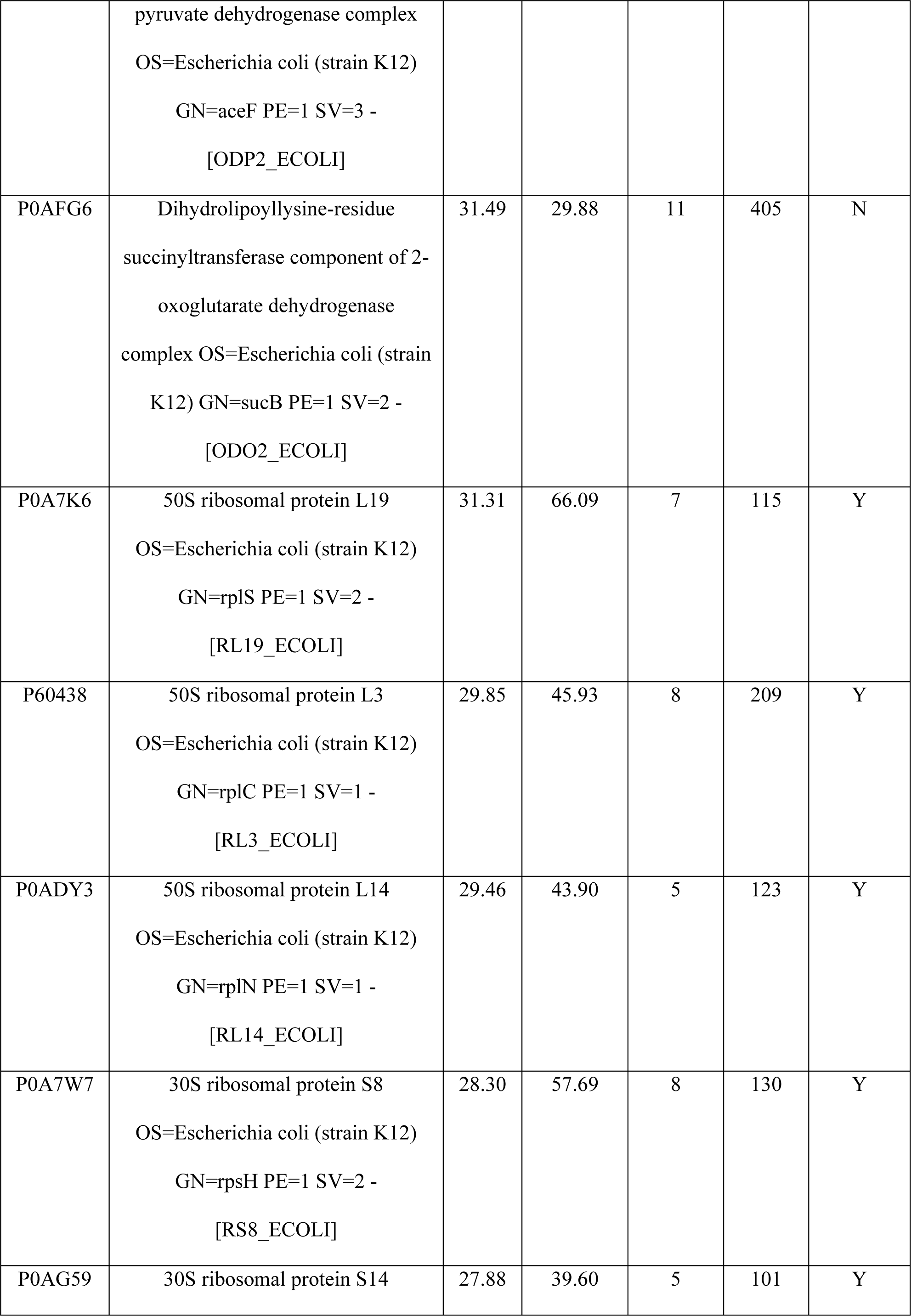
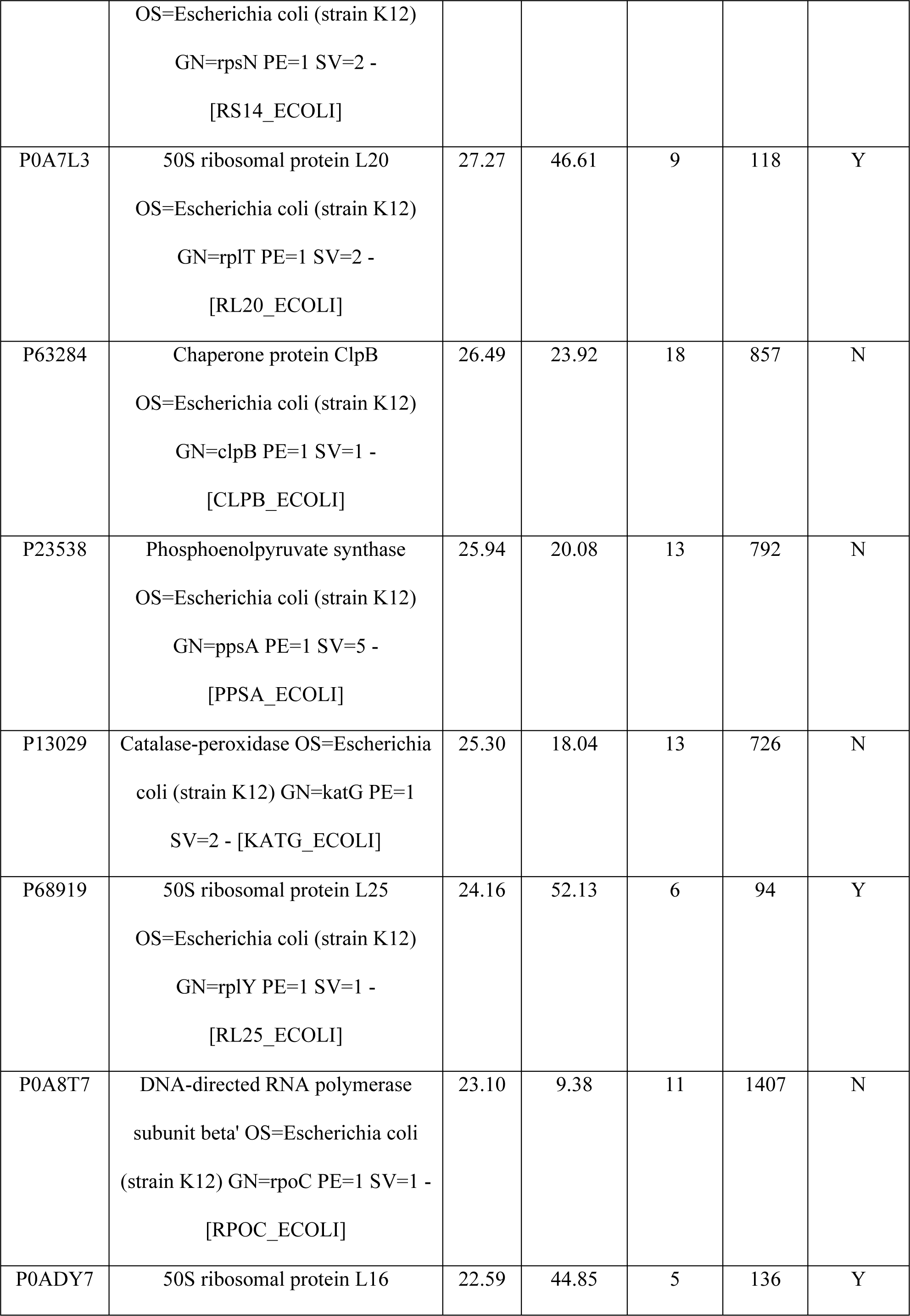
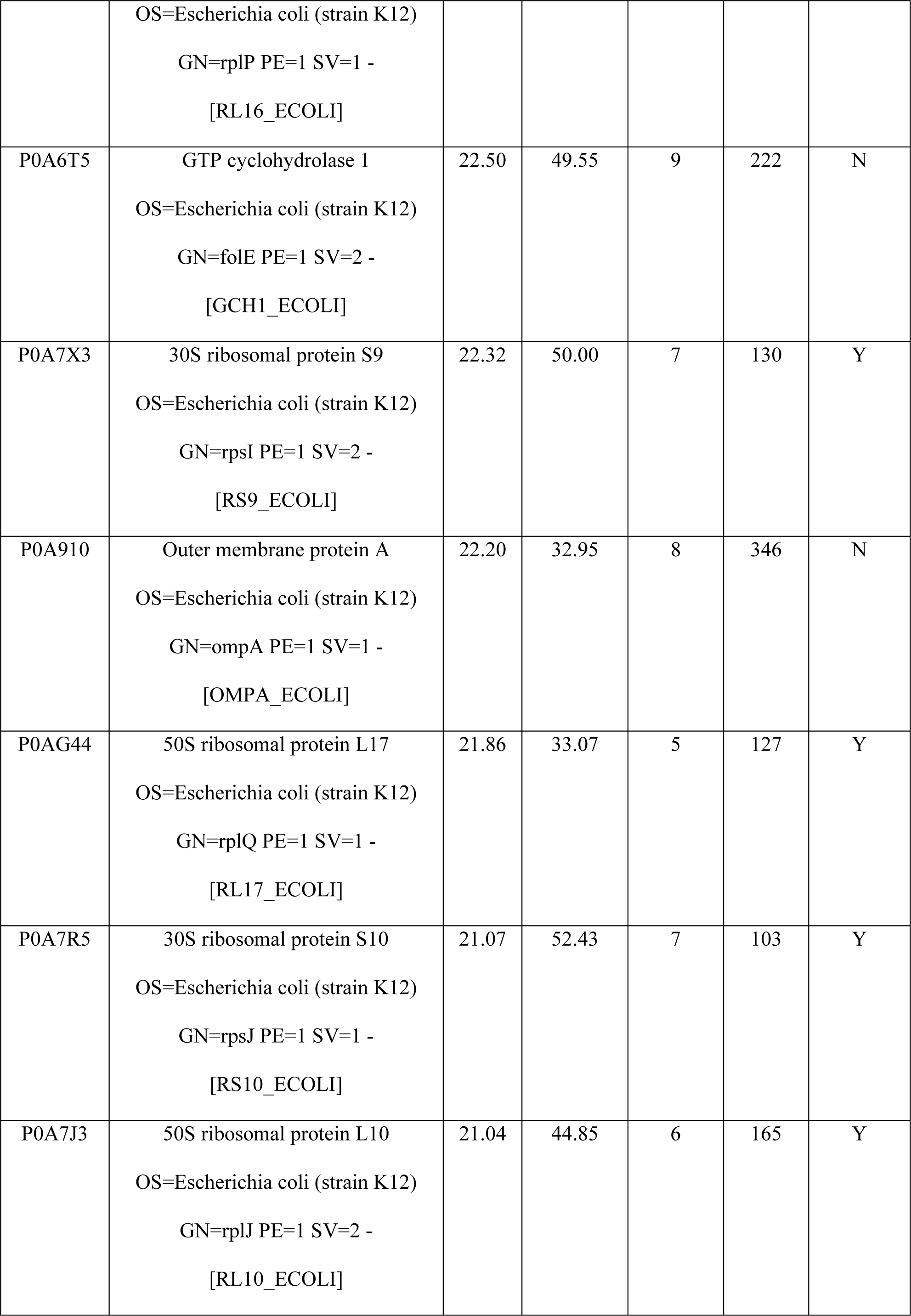
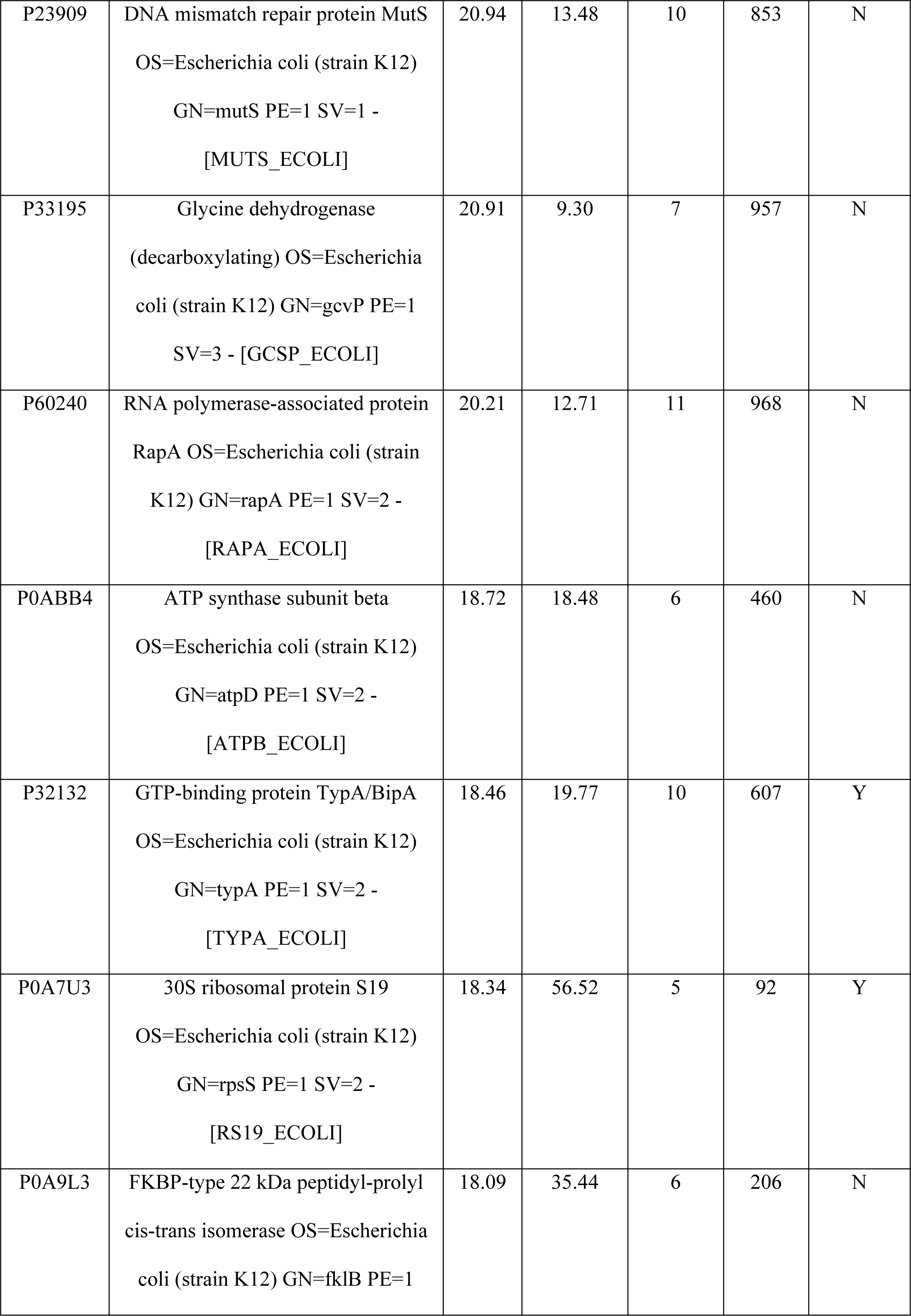
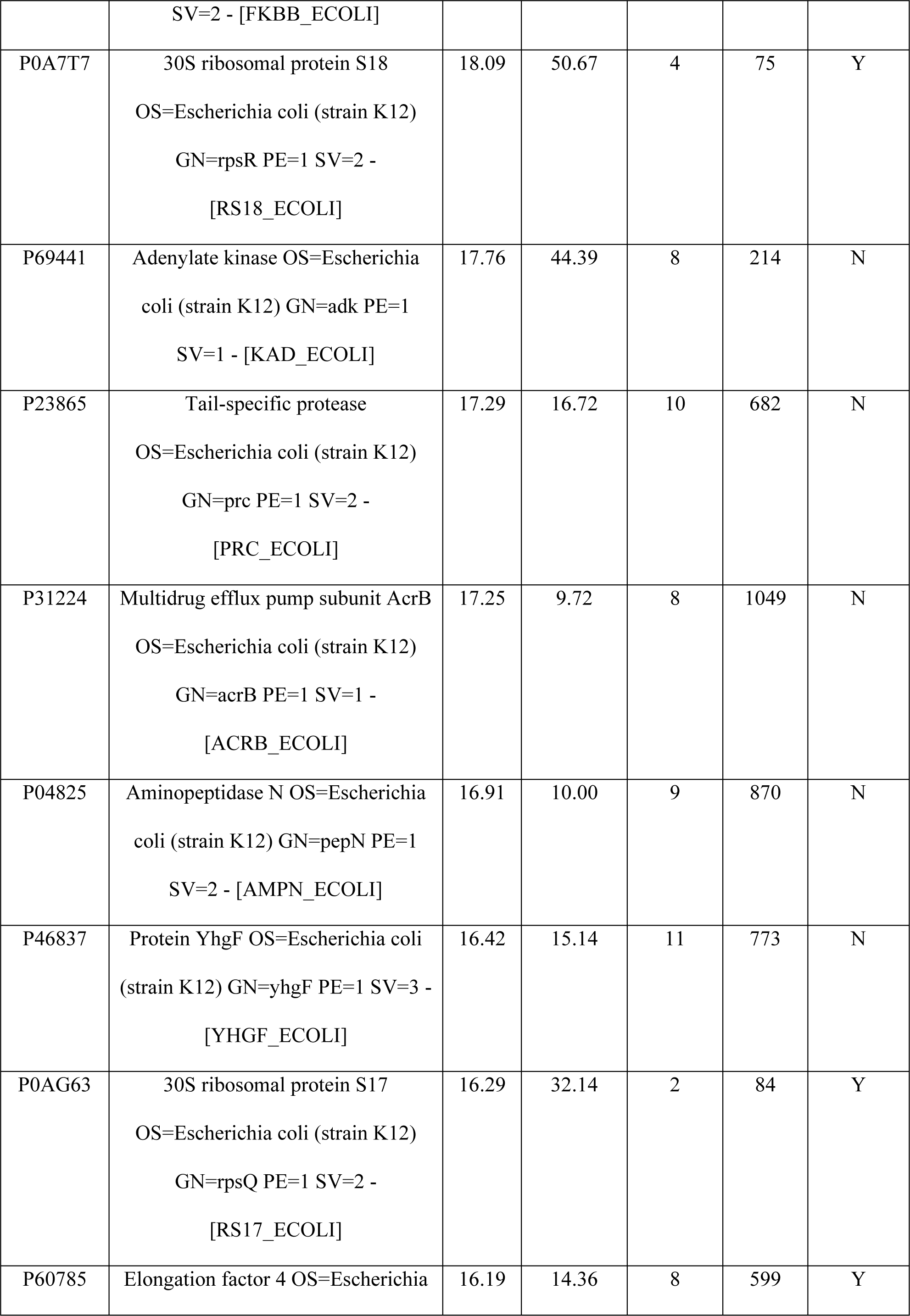
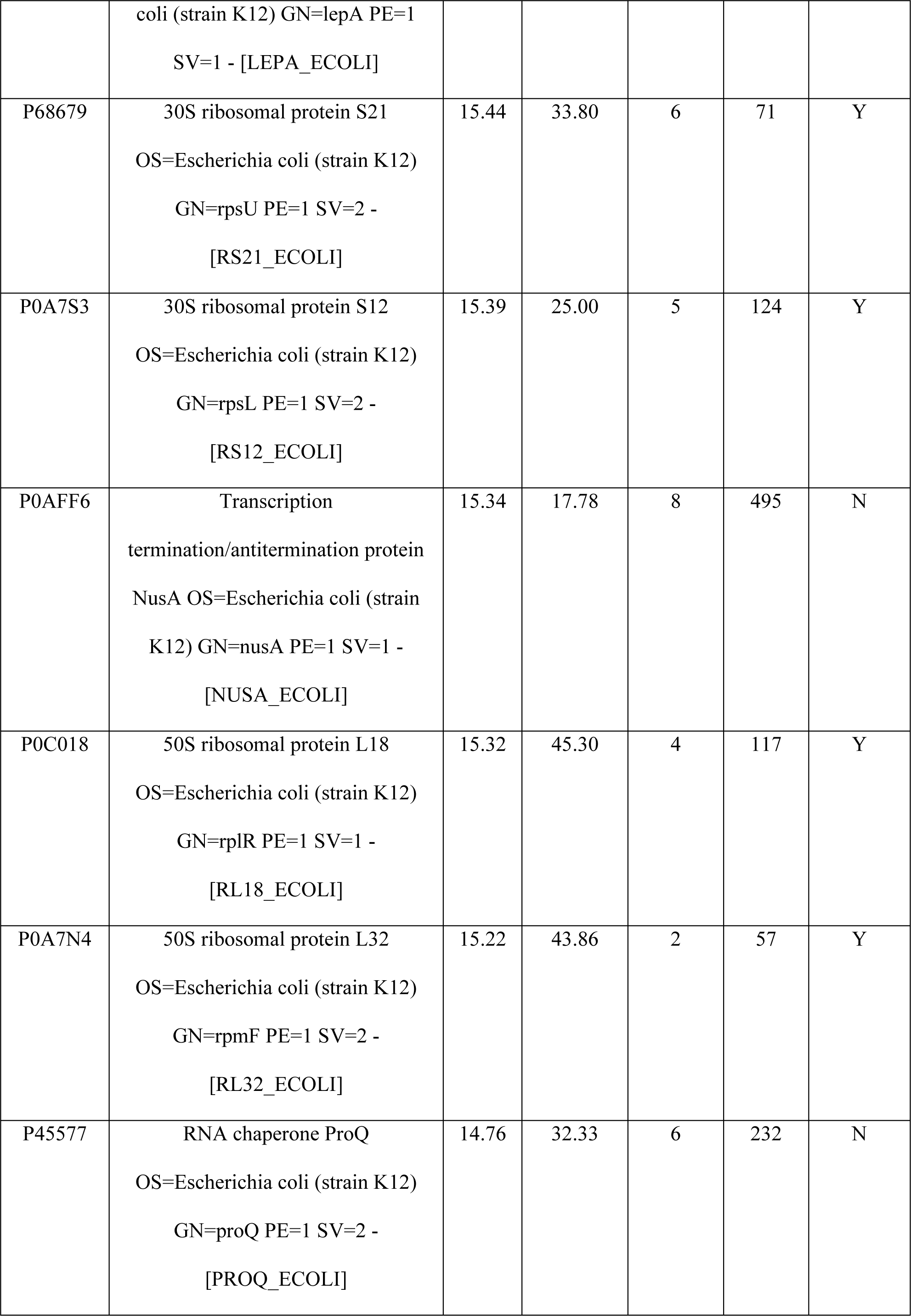
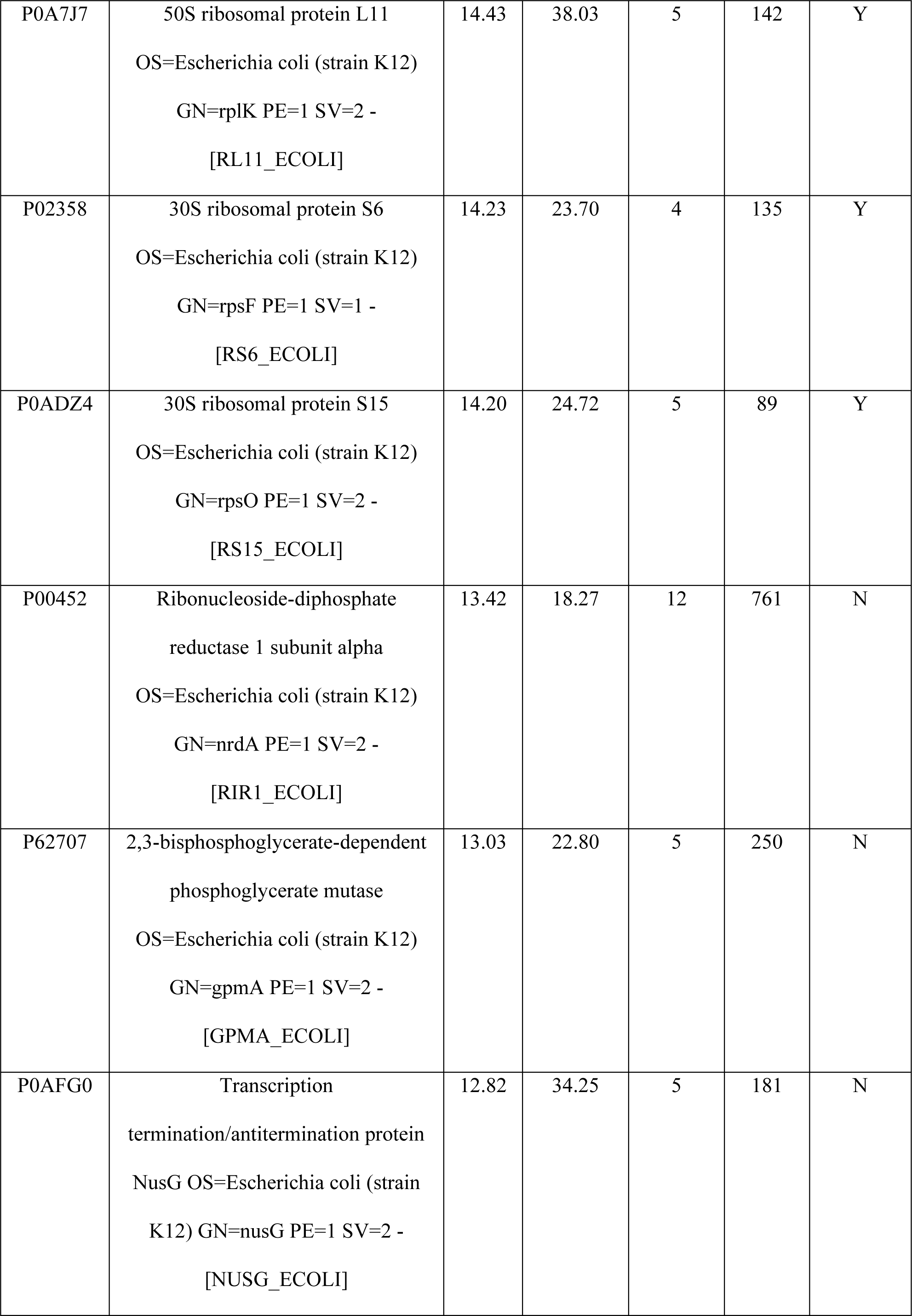
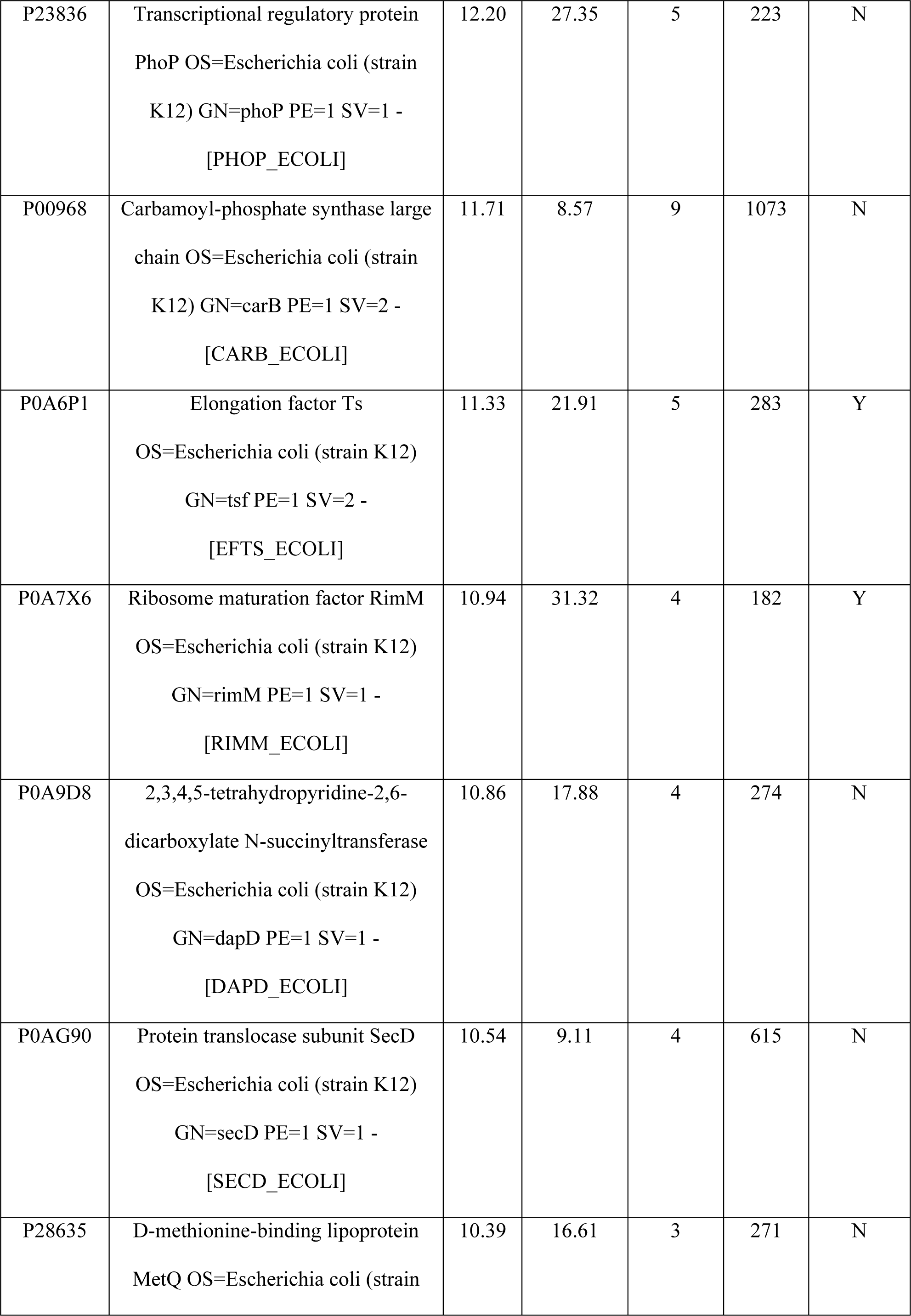
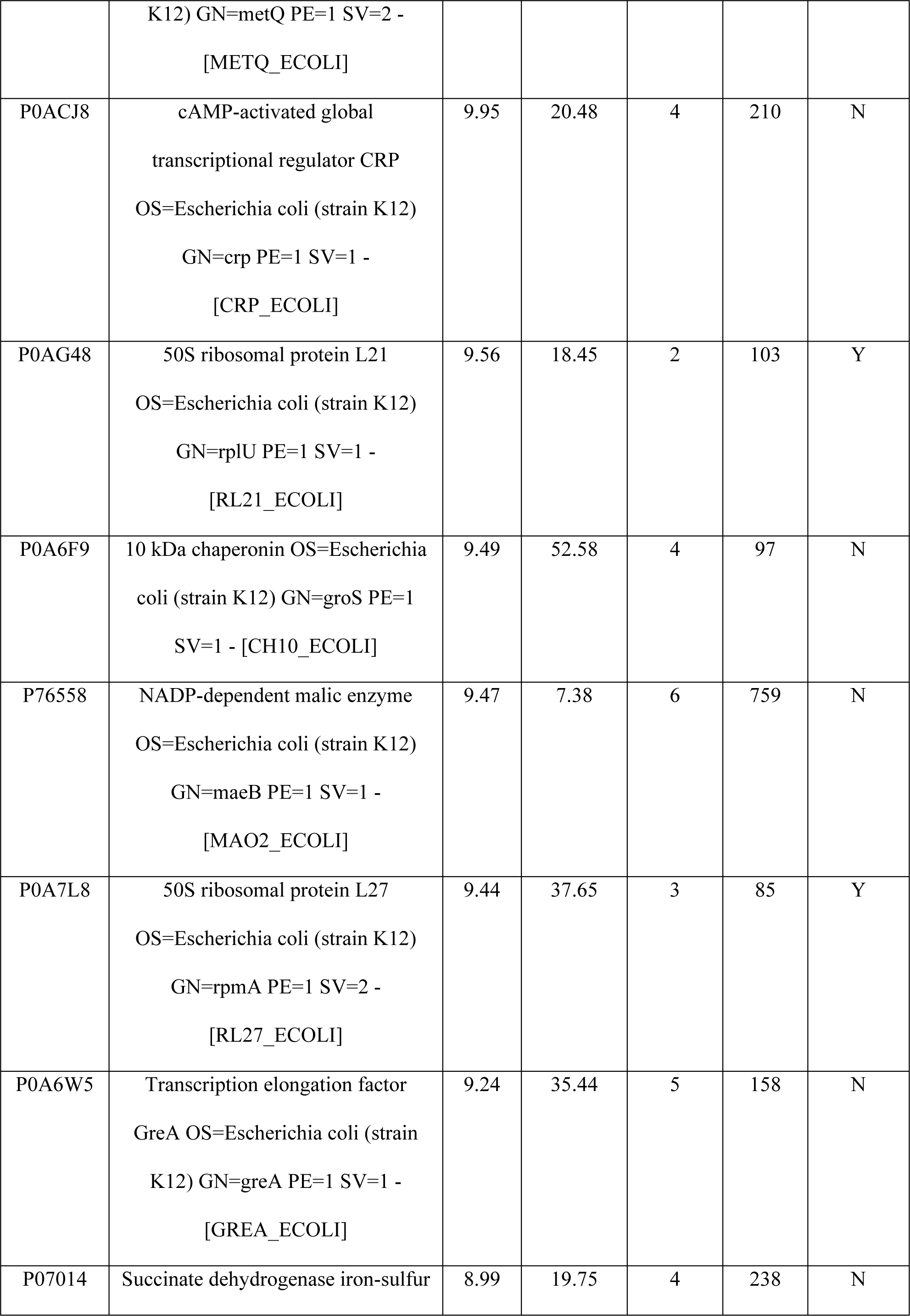
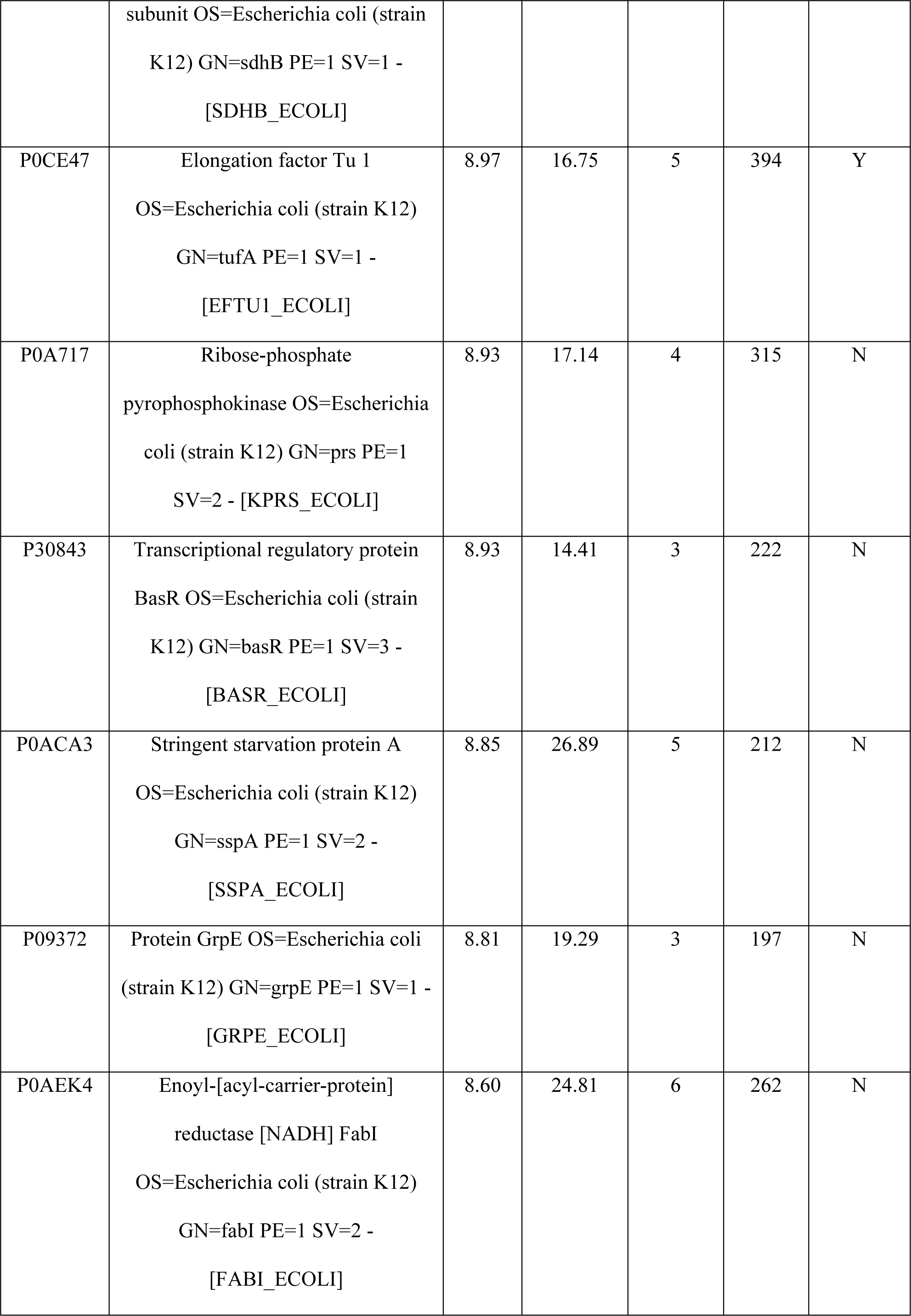
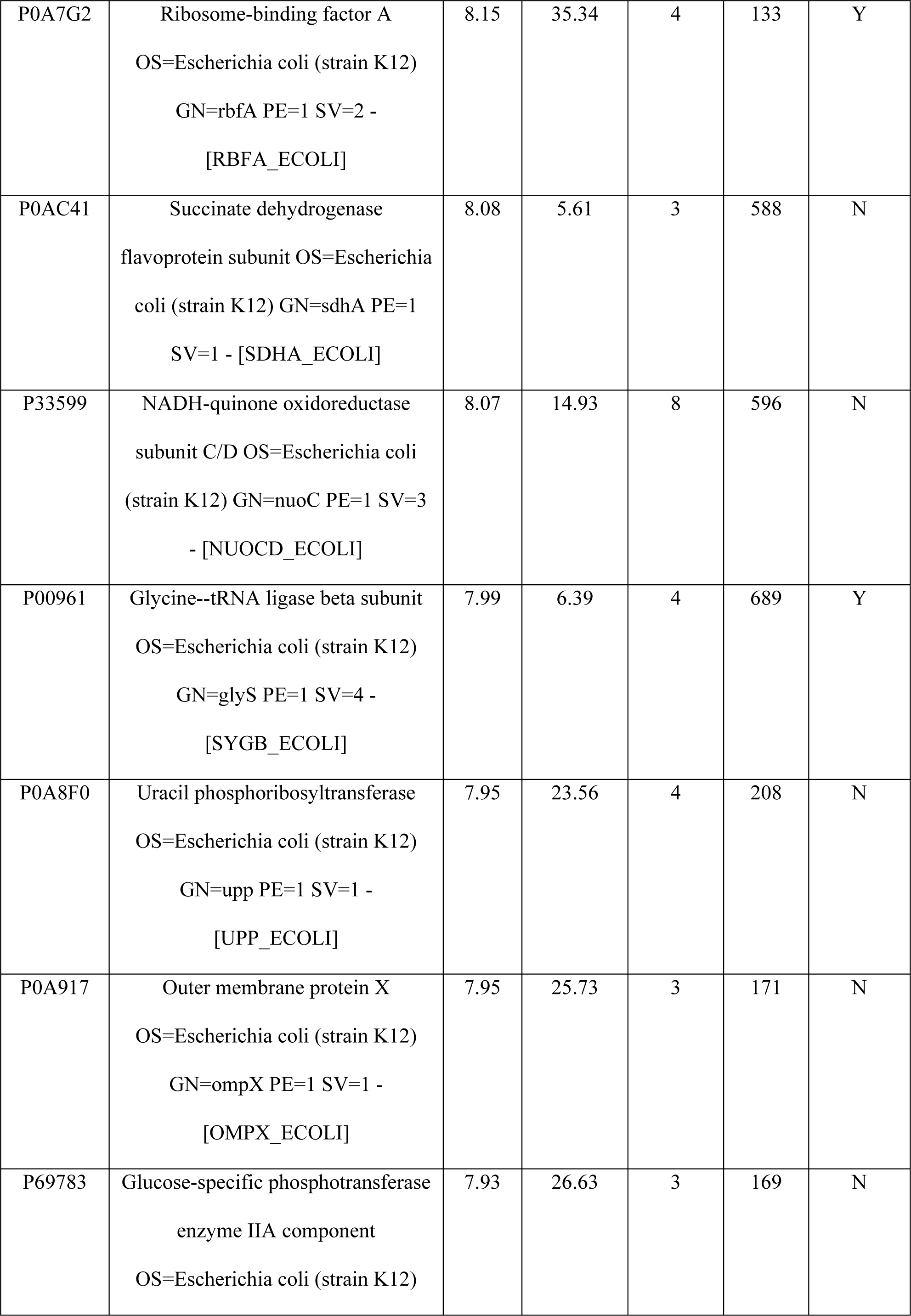
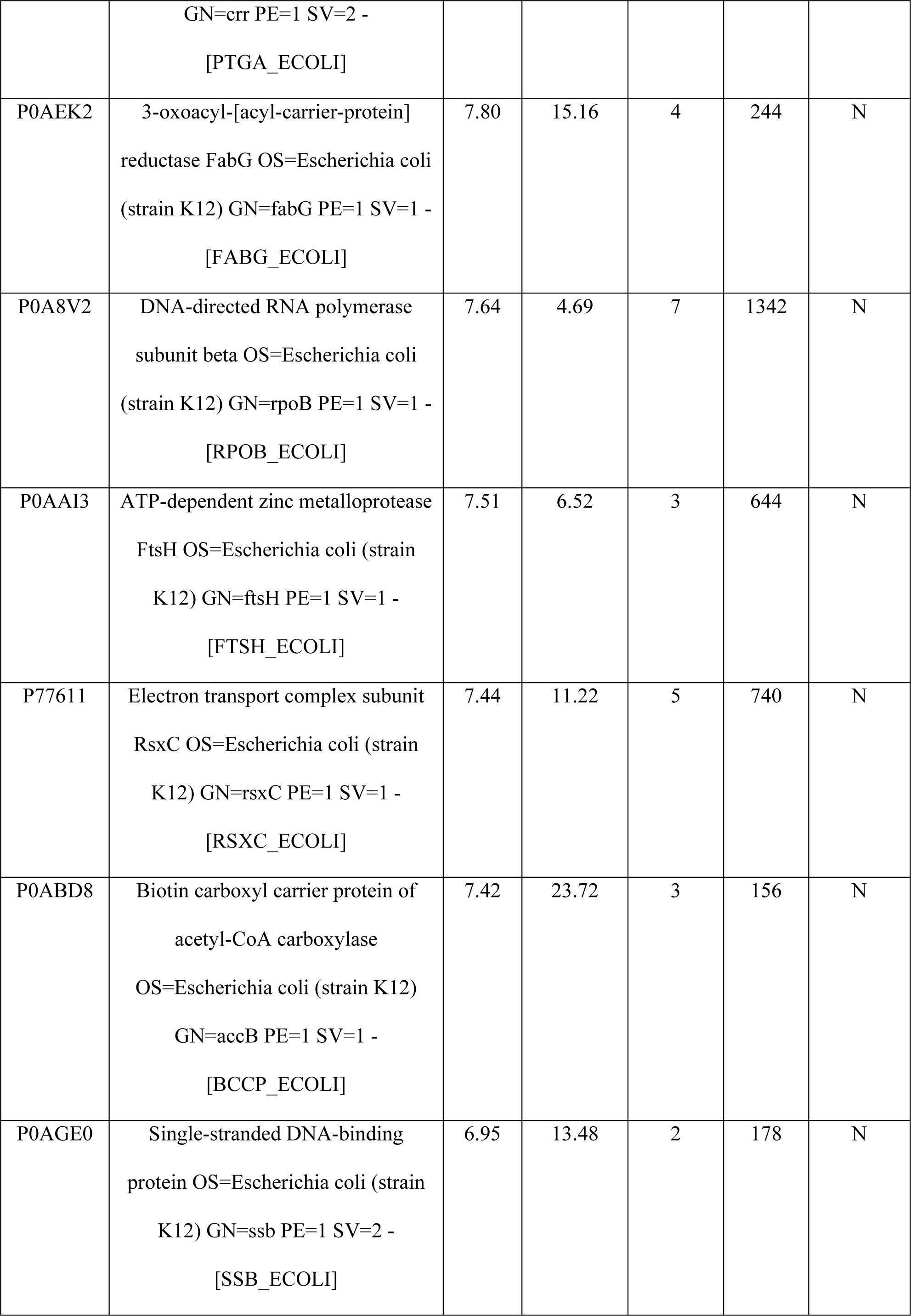
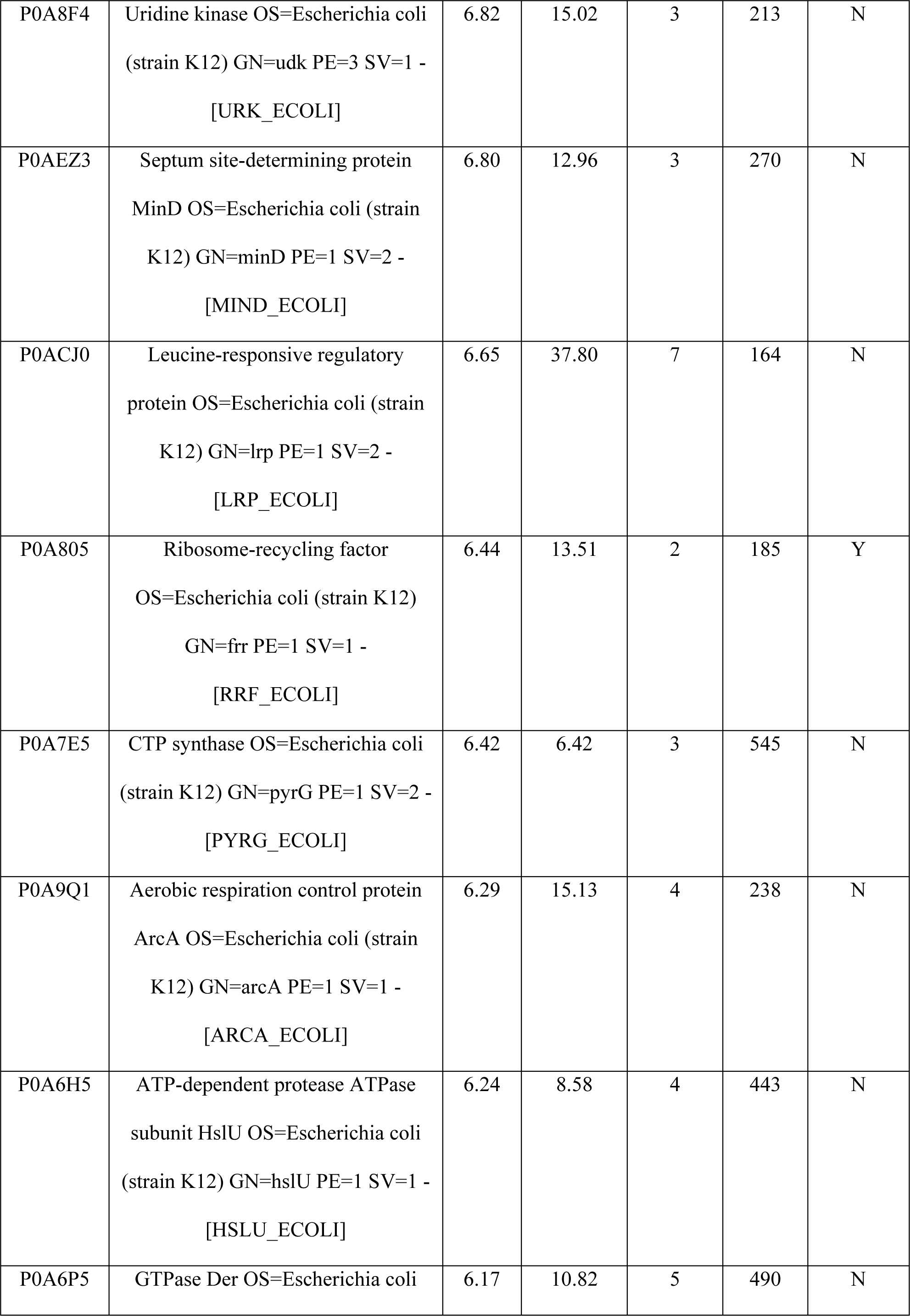
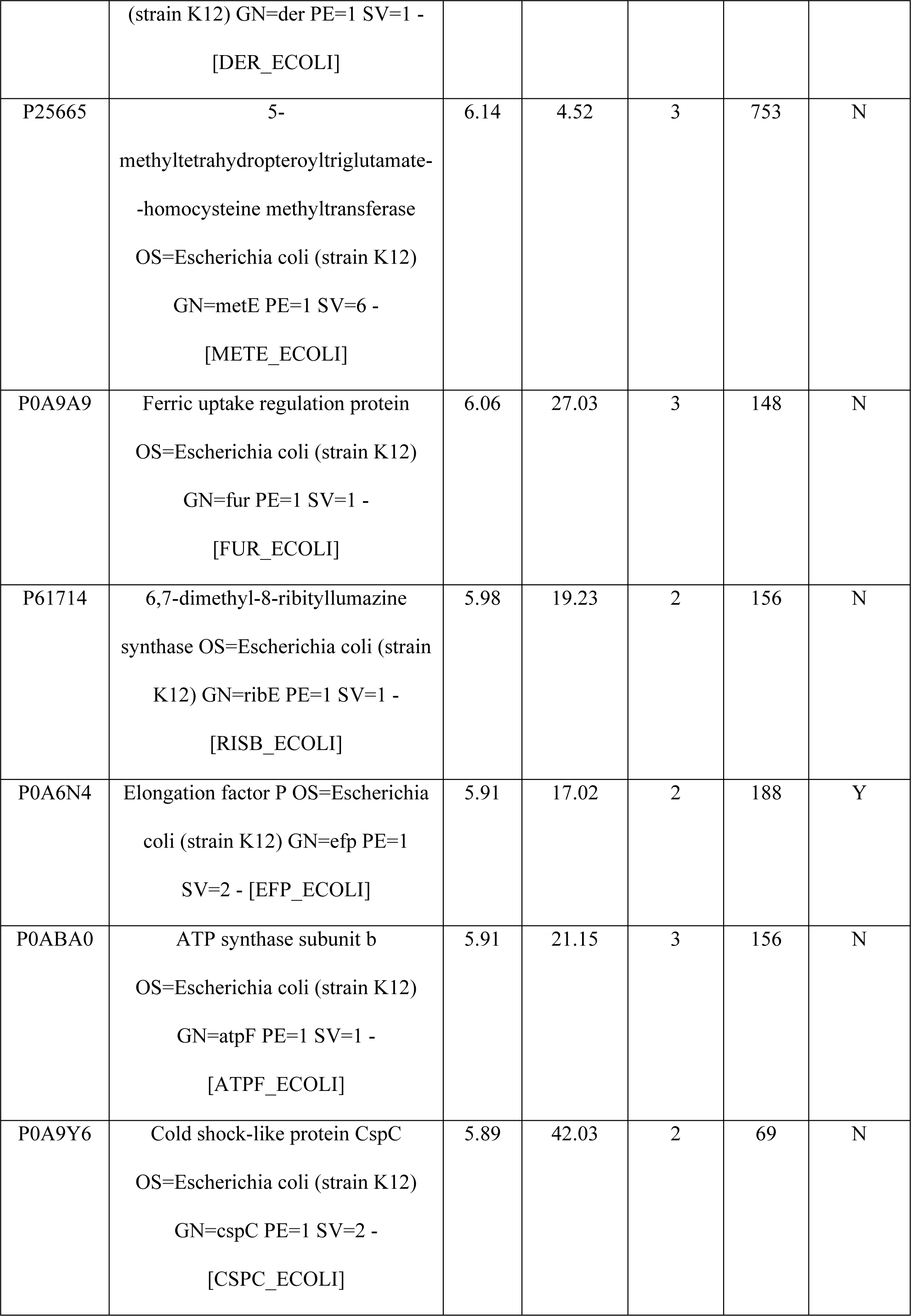
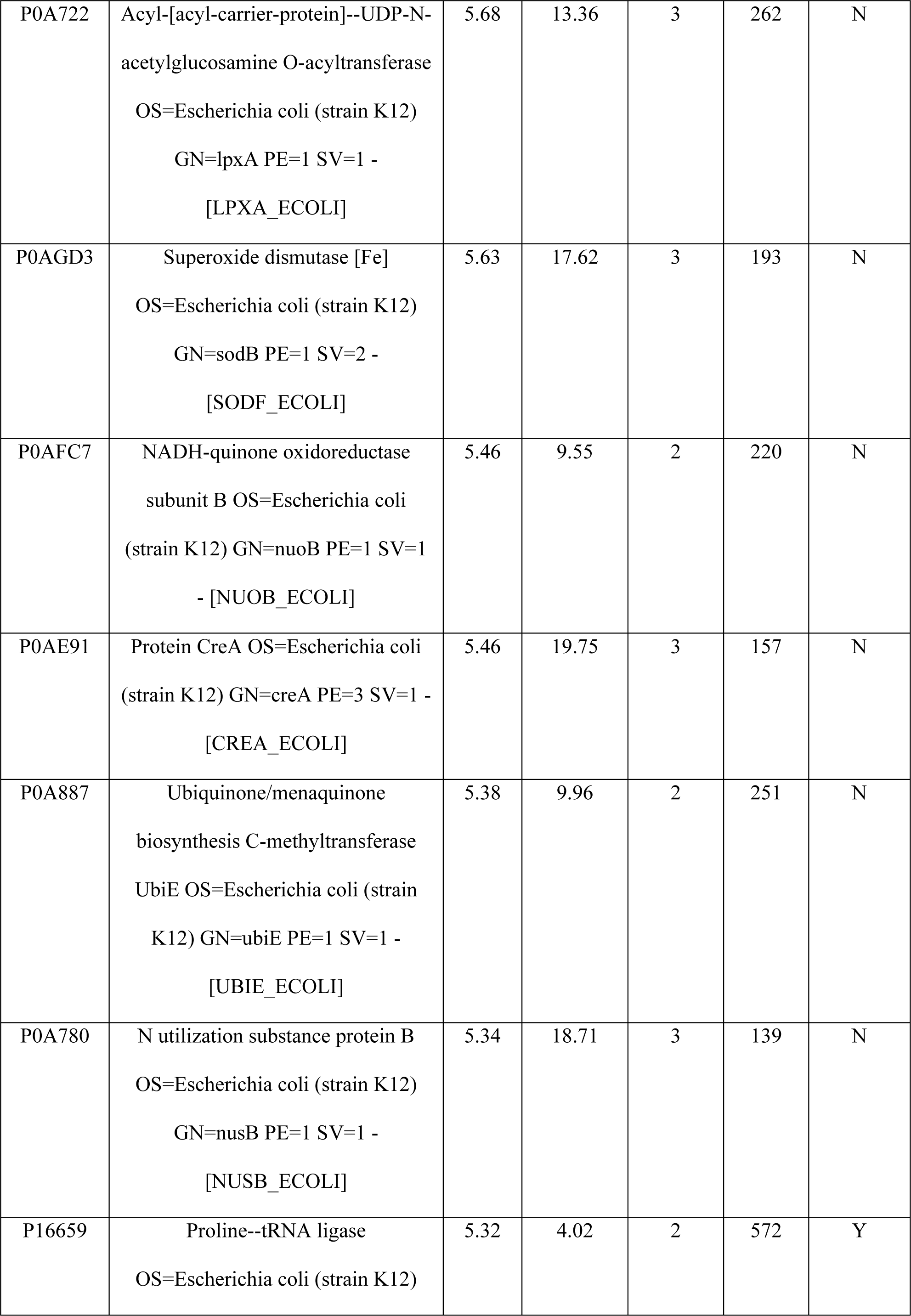
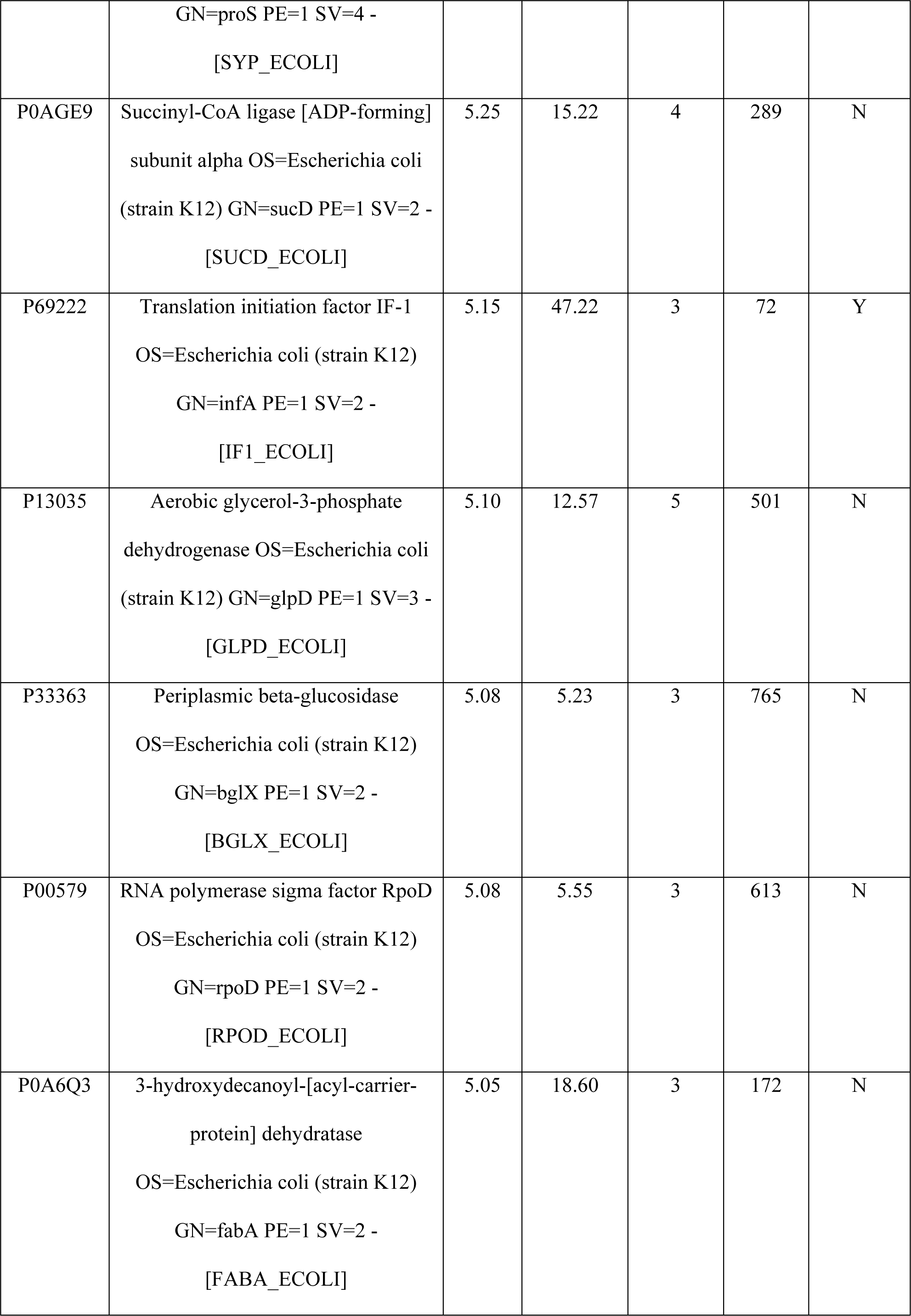
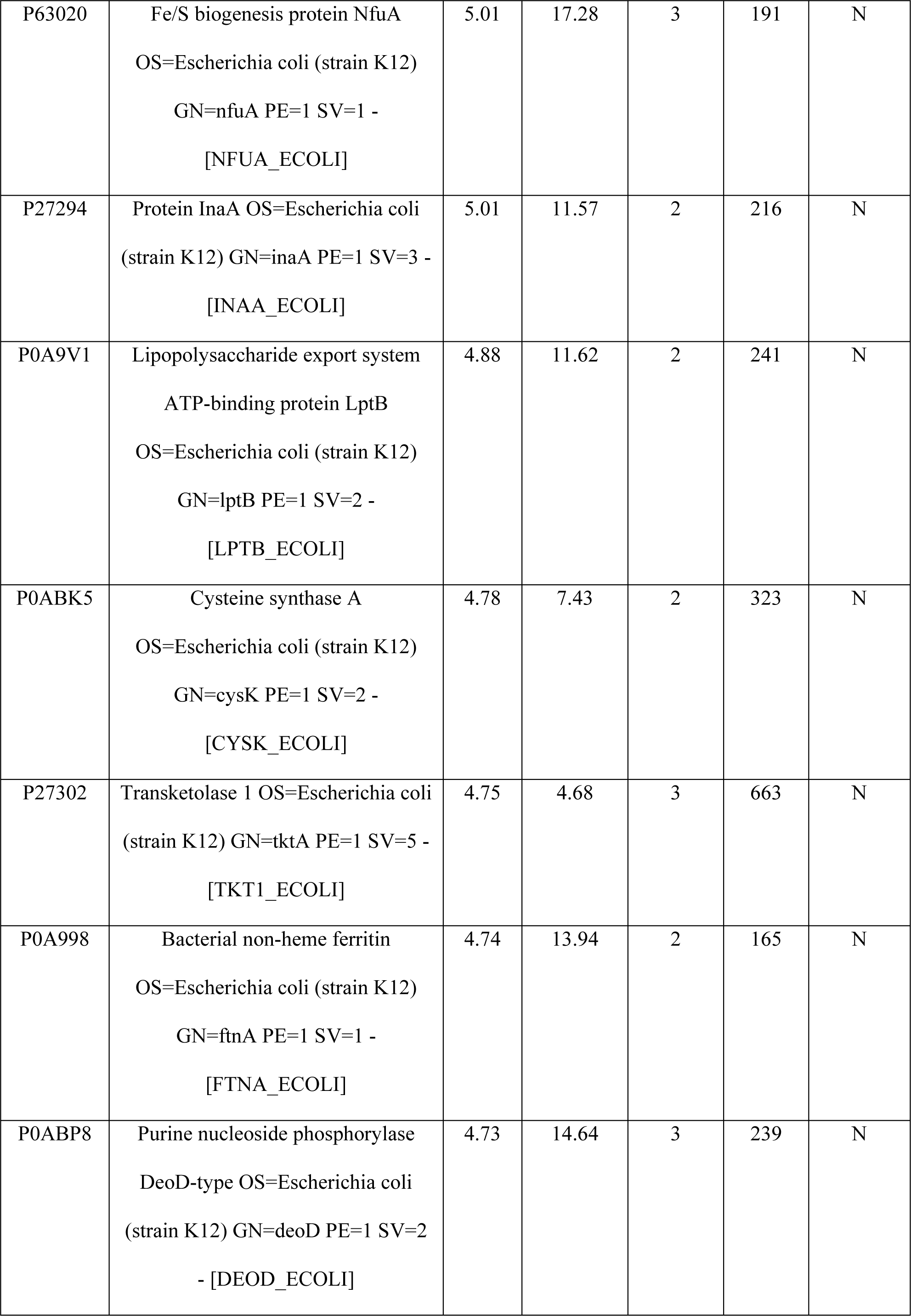
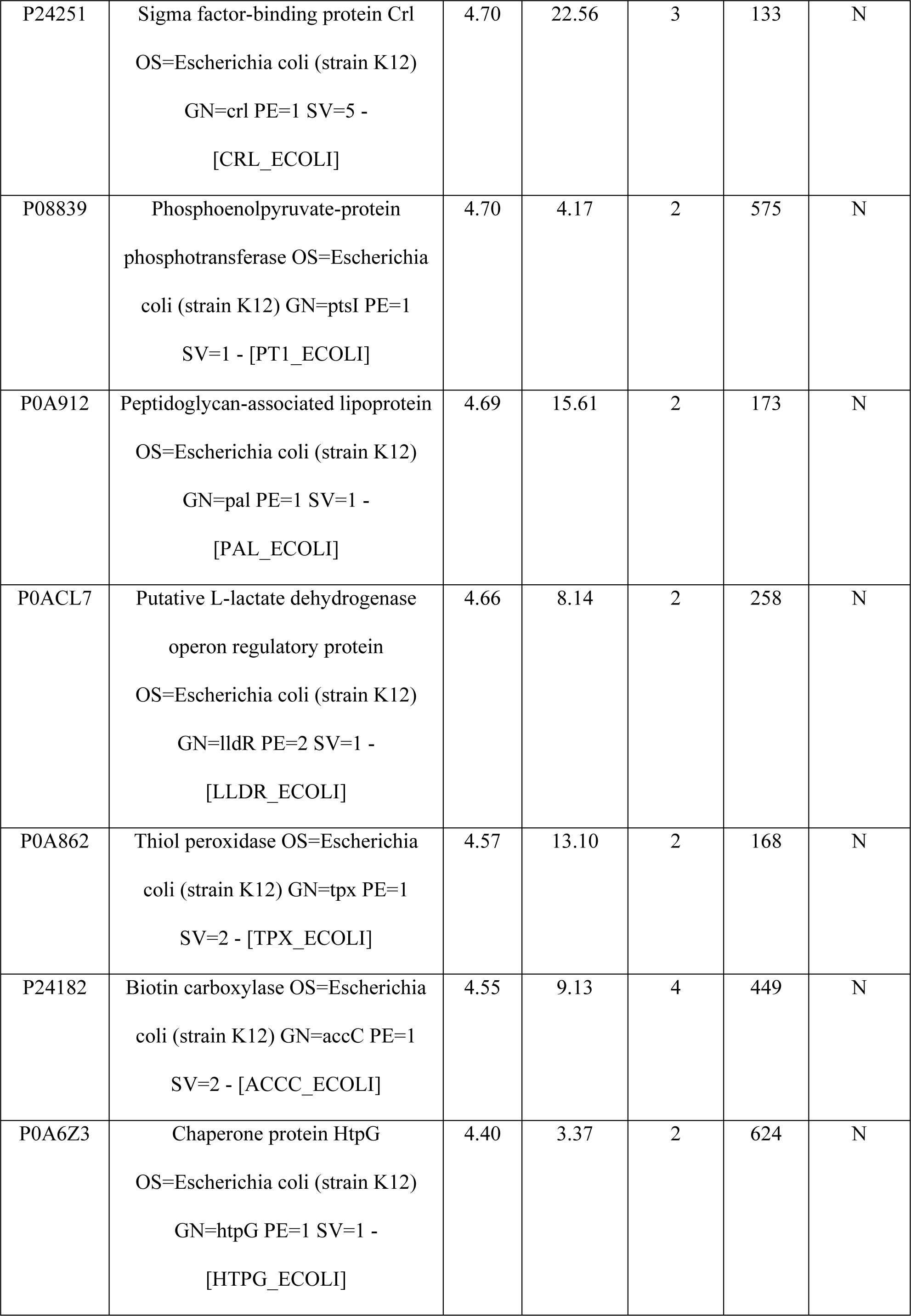
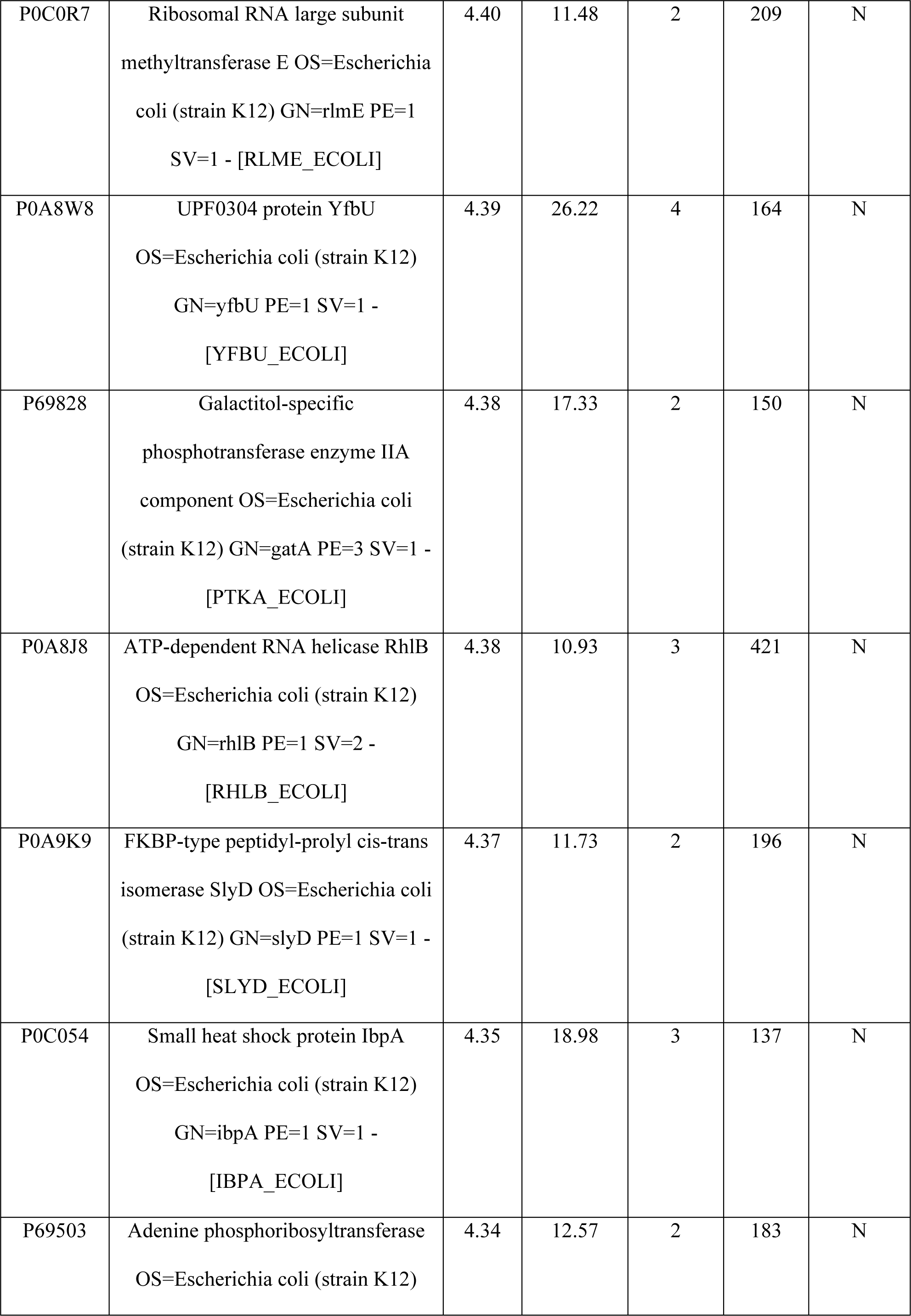
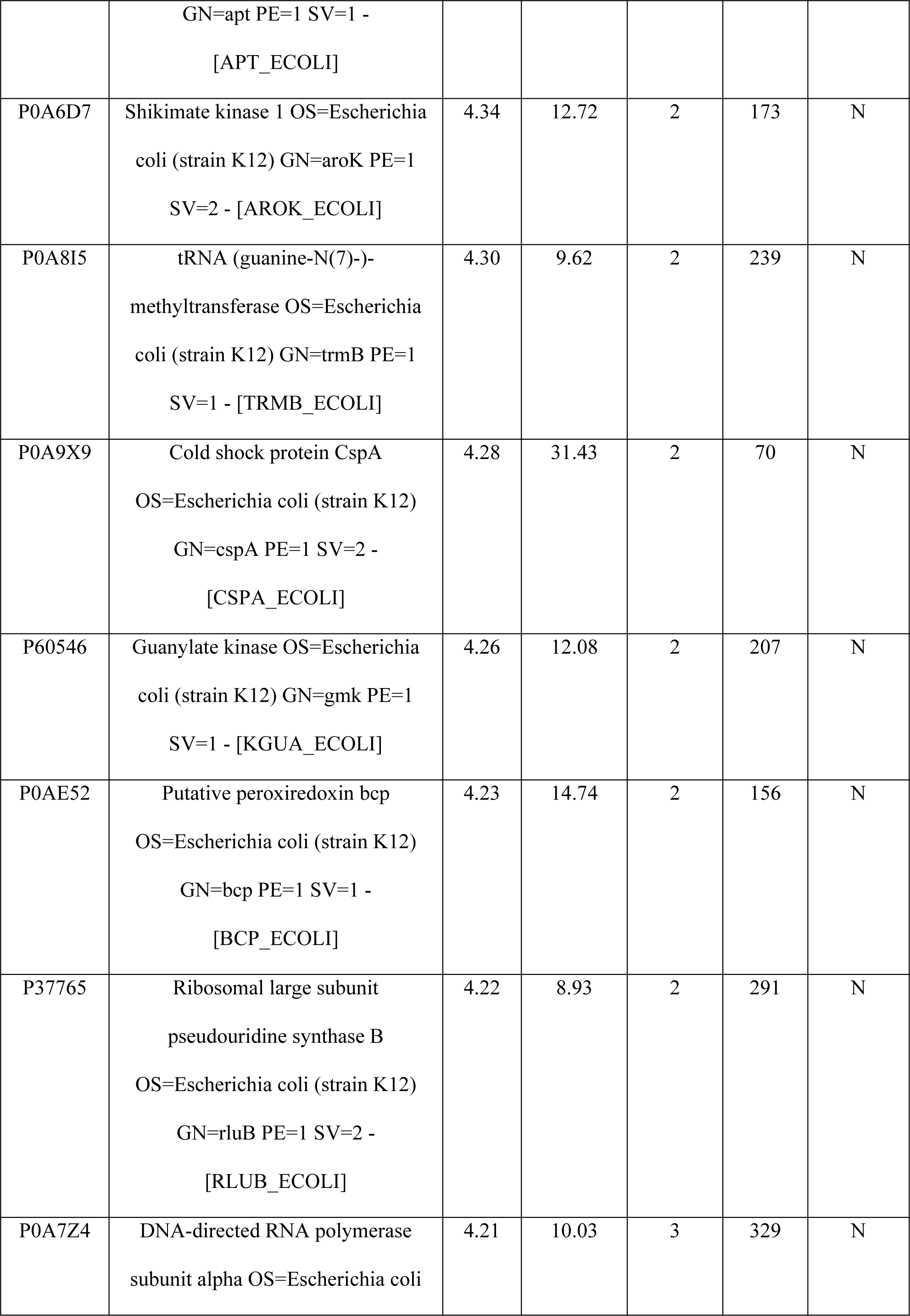
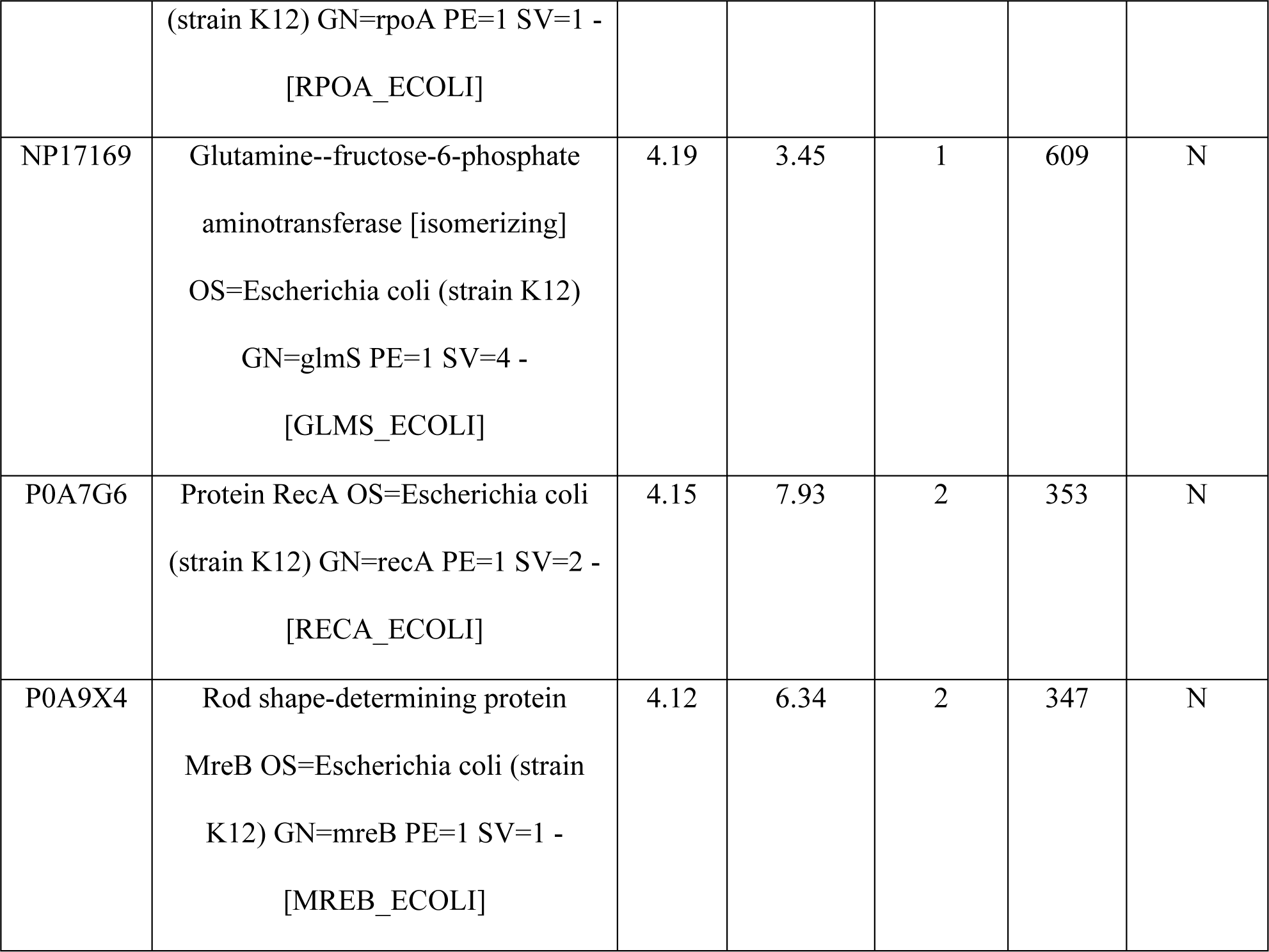
*E. coli* protein identified by LC-MS/MS that copurify with SUMO-AscA.

**Supplemental table S3.** Strains and plasmids used in this study.

| Strain | Genotype | Source |
| --- | --- | --- |
| MG1655 | *Escherichia coli* K-12 $\lambda^-$ F<sup>-</sup> | Lab stock |
| BW25113 | F<sup>-</sup> / ΔaraD-araB567 lacZ4787Δ::rrnB-3 $\lambda^-$ rph-1 ΔrhaD-rhaB568<br>hsdR514 | (23) |
| MC4100 | F<sup>-</sup> / araD139 ΔlacU169 rpsL150 thi rbsR | (49) |
| MM18 | MC4100 $\phi$(malE-lacZ) | (27) |
| BL21(DE3) | ompT hsdS<sub>B</sub> (r<sub>B</sub><sup>-</sup>, m<sub>B</sub><sup>-</sup>) gal dcm $\lambda$(DE3) | Lab stock |
| DRH959 | MG1655 ΔsecB::Kan | This study |
| DRH974 | MG1655 ΔascA | This study |
| DRH975 | MG1655 ΔascA ΔsecB::Kan | This study |
| JW3584 | BW25113 ΔsecB::Kan | (23) |
| JW1896 | BW25113 ΔascA::Kan | (23) |
| CC01 | BW25113 ΔascA (Kan<sup>S</sup>) | This study |
| EH01 | BW25113 ΔascA ΔsecB::Kan + pAscA | This study |
| DRH966 | MM18 ΔascA::Kan | This study |
| DRH969 | MM18 ΔsecB::Kan | This study |
| Plasmid | Description | Source |
| pTrc99a | High-copy number plasmid with IPTG-inducible promoter and ampicillin-resistance marker | Stratagene |
| pAscA | pTrc99a containing ascA gene inserted between *NcoI* and *BamHI* restriction sites | This study |
| pCA528 | High-copy number plasmid containing T7-inducible copy of gene encoding a His<sub>6</sub>-tagged SUMO protein from *S. cerevisiae*. | (47) |
| pCA597 | High-copy number plasmid containing T7-inducible copy of gene encoding a Strep(II)<sub>3</sub>-tagged SUMO protein from *S. cerevisiae*. | (47) |
| pDH585 | pCA528 containing *secB* gene inserted between *BsaI* and *BamHI* restriction sites | This study |
| pDH963 | pCA597 containing *ascA* gene inserted between *BsaI* and *BamHI* restriction sites | This study |
| pCS070 | pCA528 containing *ascA* gene inserted between *BsaI* and *BamHI* restriction sites | This study |
| pCS071 | pCA528 containing *ascAΔMBD* gene inserted between *BsaI* and *BamHI* restriction sites | This study |
| pCS163 | pCA597 containing *ascAΔMBD* gene inserted between *BsaI* and *BamHI* restriction sites | This study |

